# Identification of miRNA signatures for kidney renal clear cell carcinoma using the tensor-decomposition method

**DOI:** 10.1101/849679

**Authors:** Ka-Lok Ng, Y-h Taguchi

## Abstract

Cancer is a highly complex disease caused by multiple genetic factors. MicroRNA (miRNA) and mRNA expression profiles are useful for identifying prognostic biomarkers for cancer. Kidney renal clear cell carcinoma (KIRC), which accounts for more than 70% of all renal malignant tumour cases, was selected for our analysis.

Traditional methods of identifying cancer prognostic markers may not be accurate. Tensor decomposition (TD) is a useful method uncovering the underlying low-dimensional structures in the tensor. The TD-based unsupervised feature extraction method was applied to analyse mRNA and miRNA expression profiles. Biological annotations of the prognostic miRNAs and mRNAs were examined utilizing the pathway and oncogenic signature databases DIANA-miRPath and MSigDB.

TD identified the miRNA signatures and the associated genes. These genes were found to be involved in cancer-related pathways, and 23 genes were significantly correlated with the survival of KIRC patients. We demonstrated that the results are robust and not highly dependent upon the databases we selected. Compared with traditional supervised methods tested, TD achieves much better performance in selecting prognostic miRNAs and mRNAs.

These results suggest that integrated analysis using the TD-based unsupervised feature extraction technique is an effective strategy for identifying prognostic signatures in cancer studies.

## Introduction

Cancer is a highly complicated and heterogeneous disease. It is the result of a loss of cell cycle control [1], which is due to accumulation of genetic mutations, gene duplication [2], and aberrant epigenetic regulation [3, 4].Genetic mutations involving activation of proto-oncogenes to oncogenes (OCG) and inactivation of tumor-suppressing genes (TSG) may cause cancer by alternating transcription factors (TF), such as the *p53* and *ras* oncoproteins, which in turn control the expression of other genes. Gene duplication causes an elevated level of its protein product and thus favor the proliferation of cancer cells. MicroRNAs (miRNAs) are a class of small non-coding RNAs that bind to the messenger RNA (mRNA) and induce either its cleavage or impede translation repression. Several studies have indicated that abnormal miRNA expression is associated with carcinogenesis [5]. miRNAs induce cancers by acting as oncogenes (OCG) and tumor suppressor genes (TSG). An miRNA that targets the mRNA of a TSG would induce loss of the protective effect of the TSG [5, 6]. Although there have been many advancements in cancer therapy and diagnosis, many patients are unable to recover or experience recurrence after treatment. Accordingly, miRNA expression profiles are useful for identifying prognostic biomarkers for cancer diagnosis. For instance, dysregulated miRNAs were identified in urothelial carcinoma of the bladder [7]. Recent studies also suggested that miRNAs could be used as a prognostic biomarker for patients with pancreatic adenocarcinoma [8, 9]. Furthermore, by utilizing meta-analysis, it was reported that a panel of eight-miRNA signatures could serve as an effective marker for predicting overall survival in bladder cancer patients [10]. In this study, we selected kidney renal clear cell carcinoma (KIRC) for our analysis. KIRC is the most common cancer subtype of all renal malignant tumors, accounting for more than 70% of the cases (Zhang et al. 2013). Several studies have identified a few miRNA signatures that are associated with the overall survival of KIRC patients [11–13].

Typical data structures in bioinformatics are difficult to analyze because of the small number of samples with many variables. However, supervised feature extraction are effective methods for reducing the number of features. If supervised learning is applied, overfitting can occur. For example, suppose that we are seeking genes associated with aberrant expression that a disease causes. If some of those genes are also associated with gender-dependent expression while others are not, the former might be identified as less coincident with disease progression than the latter. Although gender-dependent expression is biologically acceptable, it is practically difficult to take into consideration in advance. Overlooking genes that are simultaneously associated with disease causing aberrant expression and gender-dependent expression is possible if we do not intentionally consider gender dependence. Considering labelling strictly often causes this kind of biologically unnecessary screening of genes. In contrast, unsupervised feature extraction, which is a data-driven strategy, allows us to recognize genes associated with gender-dependent aberrant expression if they are dominant, since we do not have to assume what specifically we would like to find in advance. At the same time, regularization (sparse modeling) attempts to minimize the number of features by restricting the sum of coefficients attributed to features and penalizes the use of additional variables. The disadvantage of regularization is that we must select the values of parameters that balance the prediction accuracy and the number of variables.

There are two major issues with supervised feature extraction methods: (i) class labels may not always be true, and (ii) there may be more class labels present in the dataset. As for (i), it is usual for a medical doctor to label samples by visual investigation. This sometimes results in errors, as some tumour samples can accidentally be classified as normal tissues. As for (ii), many diseases are often associated with several subclasses, e.g. cancer subtypes or different stages of disease progression. Thus, it is possible to have insufficient samples to cover all of these known subclasses. However, unsupervised methods such as principal component analysis (PCA) are often used to generate a smaller number of variables through the linear combination of original variables.

As for (i), because of the unsupervised nature as described above, mislabelling cannot generate incorrect linear combinations, since labels are used only to validate generated features, not to generate features themselves. As for (ii), again, because of the unsupervised nature, subclasses will be automatically reflected in generated features. Thus, even if we do not have enough samples to attribute to all known subclasses, features generated naturally can take these subclasses into account.

The problem with this approach is that the linear combination of many variables often prevents us from interpreting the newly generated variables. An unsupervised methodology that is suitable for the dimension reduction problems is PCA or tensor decomposition (TD)-based unsupervised feature extraction (FE) [14–27]. This method allows selection of a smaller number of variables effectively and stably. Thus, we can have a limited number of variables, which cannot be obtained by simply performing PCA and TD on a given data set.

In this paper, tensors specifically refer to mathematical objects having three or more suffices, while matrices refer to tensors with exactly two suffices. PCA is a kind of matrix factorization, while TD is a factorization method applied to a tensor. The advantages of TD over PCA is that TD needs fewer components to factorize. For example, suppose that we have 1000 variables that are formatted as either a 10 × 100 matrix or a 10 × 10 × 10 tensor. PCA applied to a matrix results in two vectors that have 10 and 100 components, respectively; thus, PCA needs in total 110 components to represent 1000 variables. In contrast, TD applied to a tensor results in three vectors, each of which has 10 components. Thus, in total, TD needs only 30 components to represent 1000 variables. Fewer components allows TD to capture features in a more efficient manner, and the results are free from overfitting the expression of 1000 variables, unlike with PCA.

## Results

Figure 1 shows the flowchart of analyses and results in this study. We applied TD-based unsupervised FE to the KIRC dataset retrieved from TCGA. It was found that 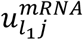 and 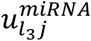 (*l*_1_ = *l*_3_*=2*) varied between the normal and tumor samples. The t-test derived P-values were 7.10 × 10^−39^ for mRNA and 2.13 × 10^−71^ for miRNA, respectively. In order to see if 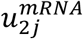 and 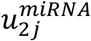 are significantly correlated, we computed the *PCC* between them, which was 0.905 (P = 1.63 × 10^−121^), indicating that they are highly correlated (Figure 2).

**Fig. 1.**
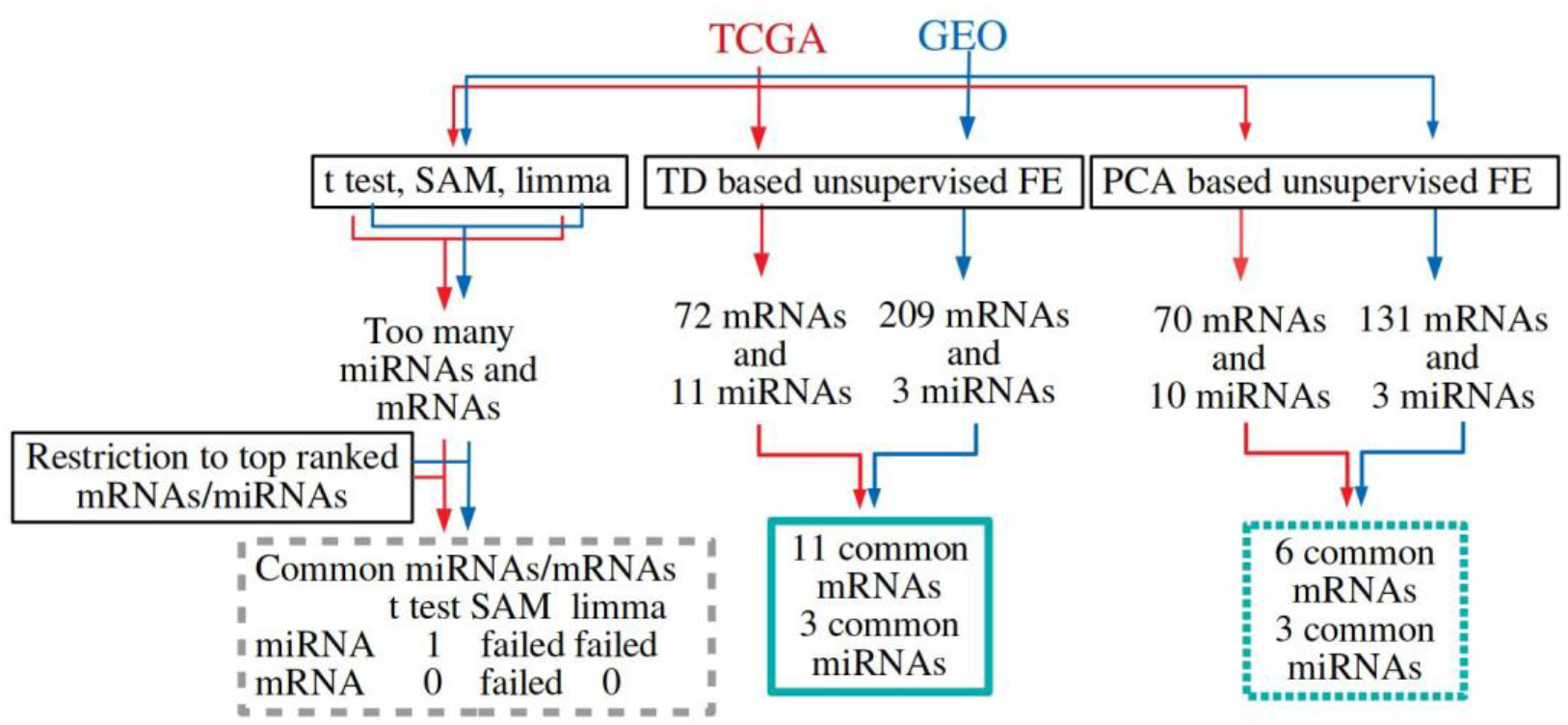
Flowchart illustrating analyses performed and the results obtained in this study. “Failed” means no successful selections from reduced number of top-ranked miRNAs or mRNAs because of assignments of *P* = 0 to too many miRNAs or mRNAs by the specified methods (for more details, see text).

**Fig. 2.**
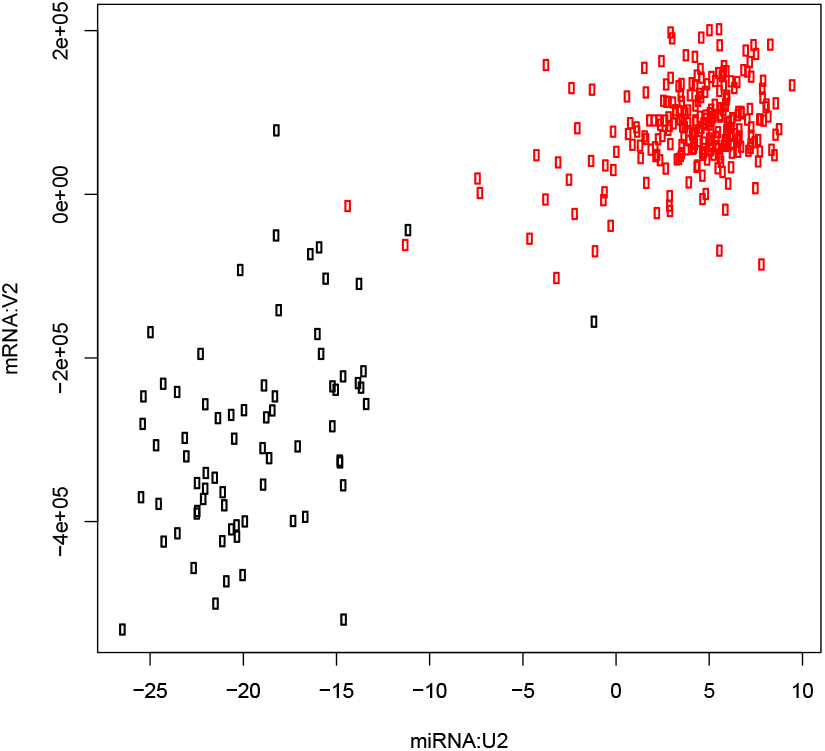
Scatter plot between 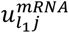 (vertical axis) and 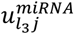 (horizontal axis). Black (red) open circle corresponds to normal (tumor) tissue.

The results of the miRNA signatures and their significant correlated genes are shown in Table 1. A total of 11 miRNAs and 72 genes were identified. To determine if these miRNAs and mRNAs are significantly correlated, we computed the *PCC* for all 11 × 72 = 792 pairs. Among them, 353 pairs were positively correlated, and 358 pairs were negatively correlated (P-values were less than 0.01 after correcting with the BH criterion). Therefore, 90% of pairs are significantly correlated. Moreover, we could successfully identify significantly correlated pairs of miRNAs and mRNAs. We noted that, among the predicted 11 miRNAs, one miRNA (miR-155) matched the result reported by Lokeshwar et al. [11].

**Table 1.**
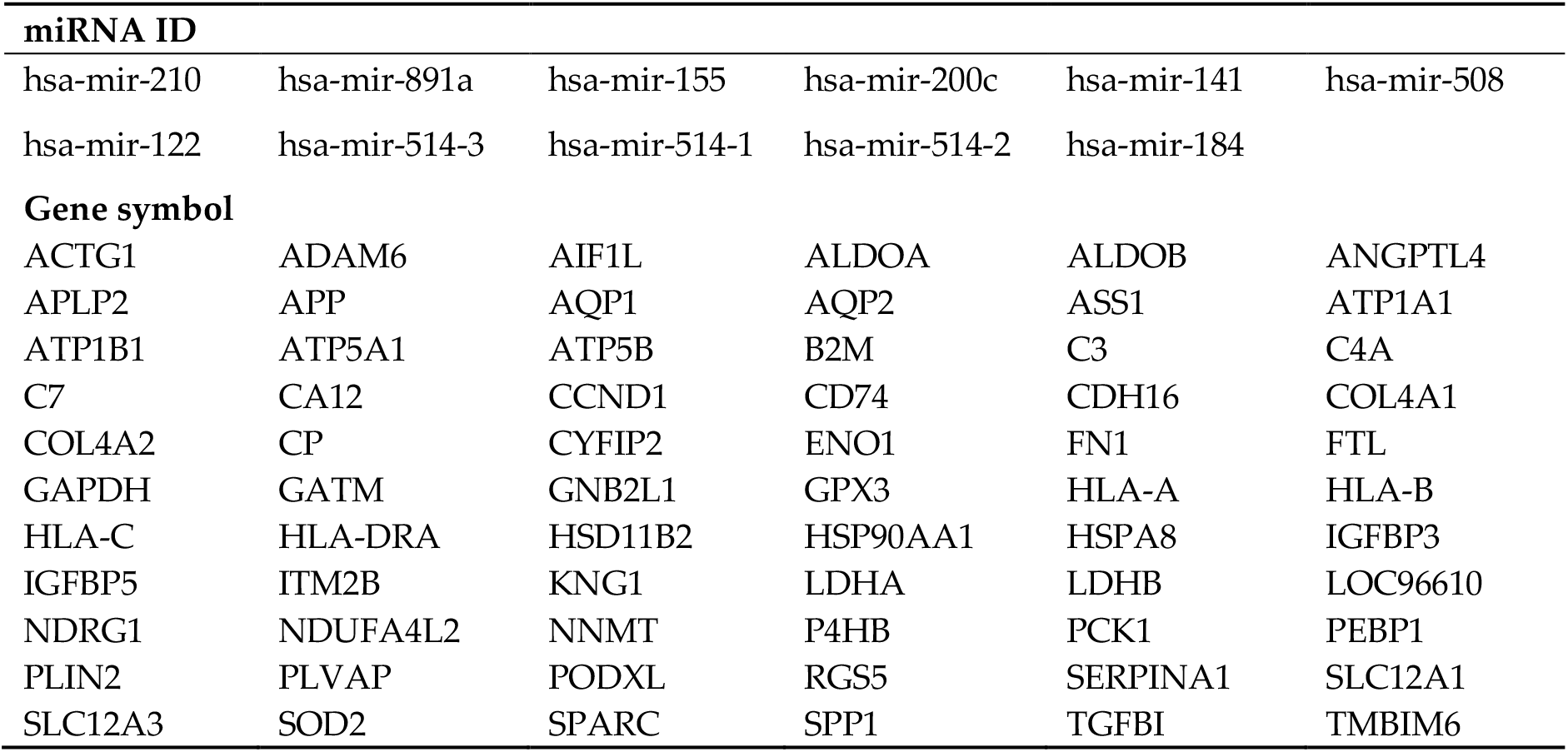
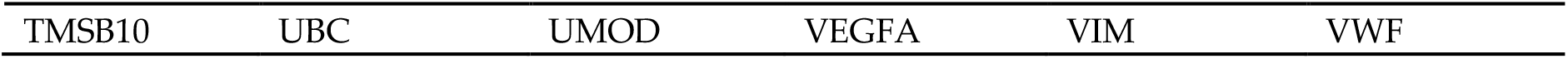
The results of the miRNA signatures and genes of KIRC patients based on the TD analysis.

Next, in order to evaluate the biological significance of selected mRNAs, we determined the top 10 oncogenic signatures of the 72 genes reported by MSigDB (Figure 3, see also Table S1). The results of the top 10 REACTOME pathways reported by MSigDB are summarized in Figure 4 (see also Table S2). These results suggest that the selected 72 mRNAs are likely related to oncogenesis. In order to further confirm if these 72 mRNAs are related to kidney cancer, we checked if these genes were linked to survival rates (Figure 5, see also Table S3). Among 72 mRNAs, 23 were significantly correlated with the survival of kidney cancer patients. This also highlights the effectiveness of our analysis. We also evaluated the identified 11 miRNAs by DIANA-mirpath. Figure 6 (see also Table S4) shows the enriched disease-related KEGG pathways (P-value < 0.05). The renal cell carcinoma pathway is identified with a significant P-value equal to 0.01613.

**Fig. 3.**
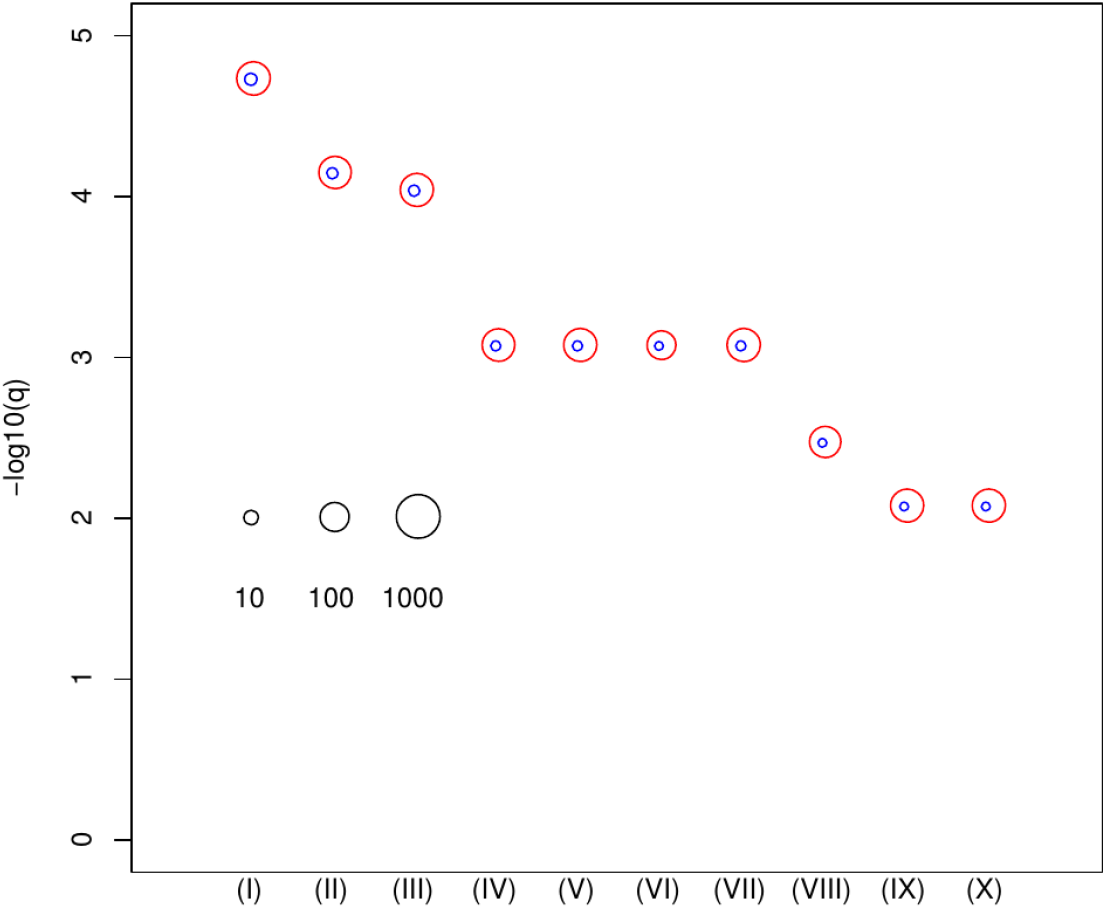
Enrichment analysis of oncogenic category in MSigDB. (I) CAMP_UP.V1_UP (II) SNF5_DN.V1_DN (III) ESC_V6.5_UP_LATE.V1_UP (IV) ESC_V6.5_UP_EARLY.V1_DN (V) ESC_J1_UP_LATE.V1_UP (VI) SIRNA_EIF4GI_UP(VII) P53_DN.V1_DN (VIII) MEL18_DN.V1_UP (IX) LTE2_UP.V1_UP (X) RPS14_DN.V1_UP. Vertical axis is the negative normal logarithmic-adjusted *P*-values. The radii of open red and blue circles show the normal logarithmic values of the number of genes in each category and those of genes included in both the category and the selected genes shown in Table 1. See Table S1 for numerical data and full descriptions.

**Fig. 4.**
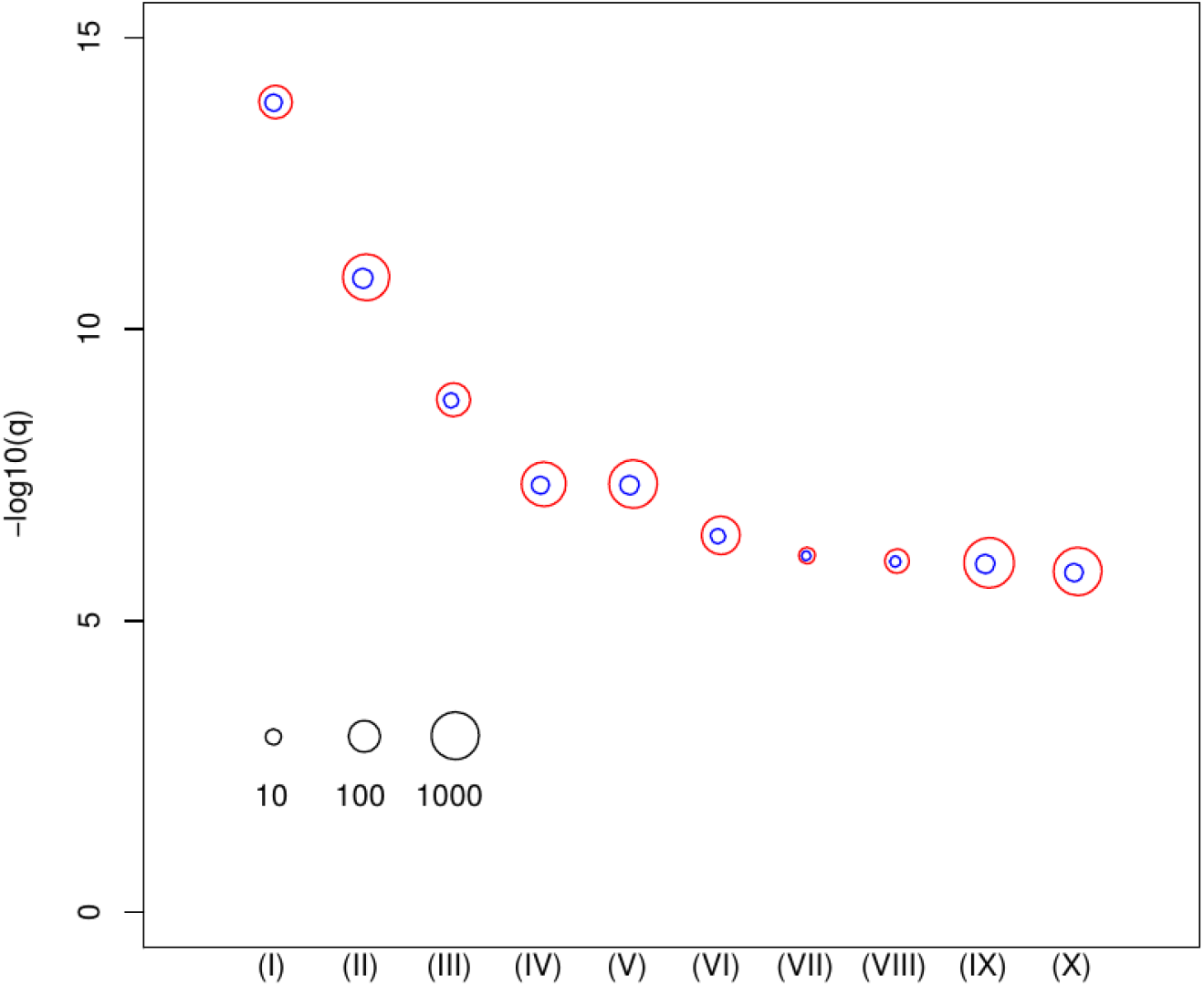
Enrichment analysis of REACTOME category in MSigDB. (I) REACTOME_regulation of insulin-like growth factor (IGF) transport and uptake by IGF binding proteins IGFBPS (II) REACTOME_cytokine signalling in immune system (III) REACTOME response to elevated platelet cytosolic CA2+ (IV) REACTOME_signalling by interleukins (V) REACTOME_innate immune system (VI) REACTOME_platelet activation, signalling, and aggregation (VII) REACTOME_endosomal vacuolar pathway (VIII) REACTOME_gloconeogenesis (IX) REACTOME_post-translational protein modification (X) REACTOME_disease. The radii of open red and blue circles show the normal logarithmic values of the number of genes in each category and those of genes included in both the category and the selected genes shown in Table 1. See Table S2 for numerical data and full descriptions.

**Fig. 5.**
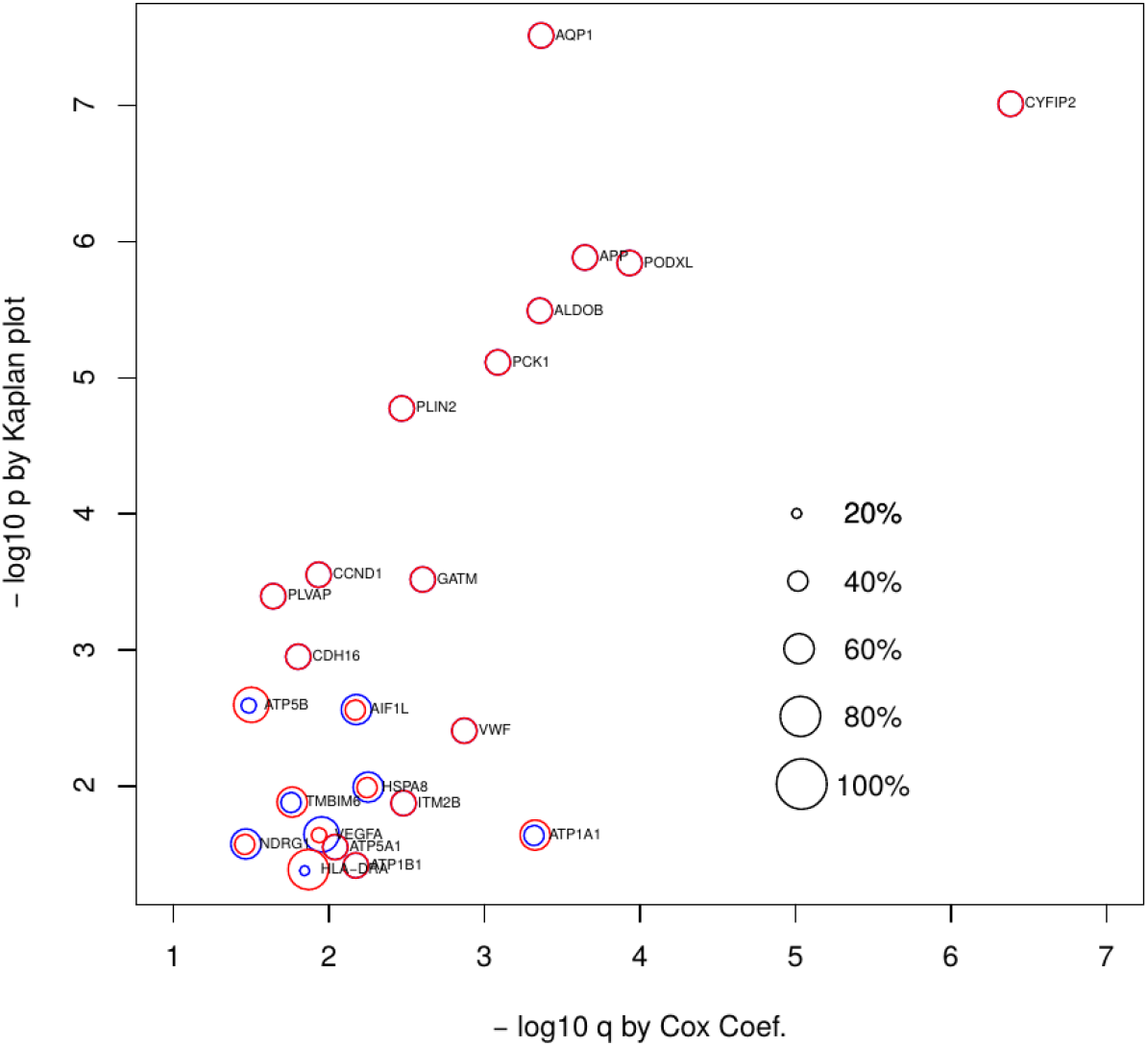
Survival analysis of 24 genes from Table 1 that significantly contribute to patients’ survival. Vertical axis: negative normal logarithmic values of *P*-values computed by Kaplan plot. Horizontal axis: negative normal logarithmic values of adjusted *P*-values computed by Cox analysis. Red open circles indicate lower expression percentile of patient groups. Only when they are not 50%, upper expression percentiles are displayed with blue circles. See Table S3 for numerical data and full descriptions.

**Fig. 6.**
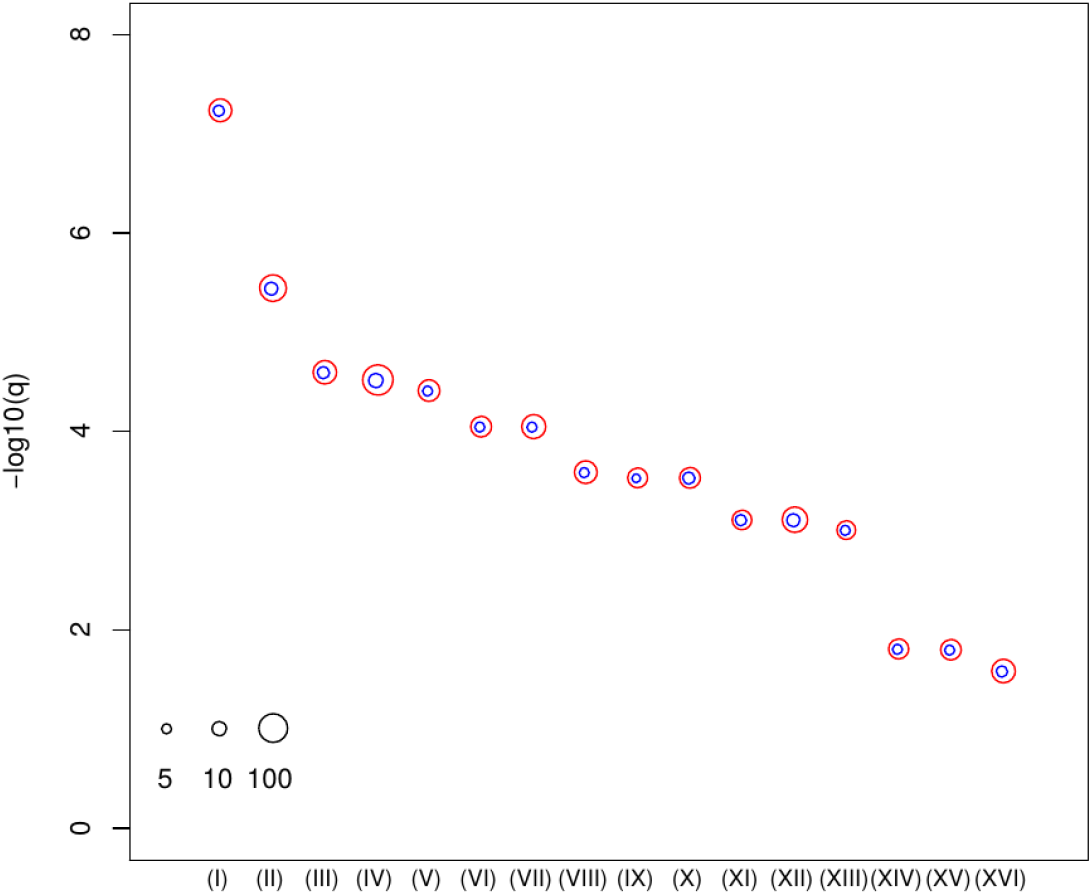
Enrichment analysis of KEGG pathway provided by DIANA-mirpath to which miRNAs in Table 1 were uploaded. Vertical axis: negative normal logarithmic values of adjusted *P*-values. (I) Chronic myeloid leukemia (II) Proteoglycans in cancer (III) Prostate cancer (IV) Pathways in cancer (V) Pancreatic cancer (VI) Glioma (VII) Hepatitis B (VIII) Small cell lung cancer (IX) Non-small cell lung cancer (X) Colorectal cancer (XI) Endometrial cancer (XII) Viral carcinogenesis (XIII) Bladder cancer (XIV) Melanoma (XV) Renal cell carcinoma (XVI) Hepatitis C. The radii of red open circles indicate the normal logarithmic values of the number of genes in each category targeted by miRNAs in Table 1 whose normal logarithmic numbers are proportional to the radii of blue open circles. See Table S4 for numerical values and full description.

The top signature in Table S1 [Fig. 3(I)] is related to the cAMP signaling pathway. Targeting the cAMP pathway is an effective treatment for kidney cancer [28, 29]. The second signature in Table S1 [Fig. 3(II)] is the *Snf5* gene expression profile of a murine model (Mouse Embryonic Fibroblast (MEF) cells) that closely resembles that of human SNF5-deficient rhabdoid tumors (pediatric soft tissue sarcoma that arises in the kidney, the liver, and the peripheral nerves) [30]. Impairment of the SWI/SNF chromatin remodeling complex plays an important role in the development and aggressiveness of clear cell renal cell carcinoma [31]. The sixth signature in Table S1 [Fig. 3 (VI)] comes from a study of the effects of knockdown of the gene family of eukaryotic translation initiation factors (EIF) by RNAi in MCF10A cells. EIF3b is a promising prognostic biomarker and a potential therapeutic target for patients with clear cell renal cell carcinoma [32], and EIF4GI is a target for cancer therapeutics [33].

The top pathway in Table S2 [Fig. 4 (I)] is the ‘Pathway of regulation of IGF activity by IGFBP’. Studies show that insulin-like growth factors (IGFs) and insulin play a stimulatory role for renal cancer cells [34, 35]. Patients with IGF-1 receptor overexpression have a 70% increased risk of death [36]. Moreover, this overexpression has been shown to increase kidney cancer risk in middle-aged male smokers [37]. The second pathway in Table S2 [Fig. 4(II)] is ‘Cytokine Signaling in Immune system’. Cytokines are important biomolecules that play essential roles in tumor formation [38], and they are therapeutic targets [39, 40]. The IL-6 cytokine family can serve as useful diagnostic and prognostic biomarkers. In fact, IL-6 is a potential target in cancer therapy [41, 42]. Ishibashi et al. reported that IL-6 suppresses the expression of the cytokine signaling-3 (*SOCS3*) gene and is associated with poor prognosis of kidney cancer patients [43].

Table S3 [Fig. 5] shows the significant relationships between the predicted 23 mRNAs and the patients’ survival rates. For some of the 23 genes, patients cannot be divided equally based on expression of considered genes in order to get significant *P*-values for the Kaplan-Meier plots. A majority of the mRNAs (15 out of 23) are associated with P-values less than 0.05, with 50/50 divisions based on the level of gene expression. Among the 16 KEGG pathways predicted by DIANA-mirpath (Table S4 and Fig. 6), 14 are directly related to cancers, except for Hepatitis B and Hepatitis C. Therefore, we correctly identified miRNA signatures that are cancer-related.

In order to validate the robustness of our findings, we employed an independent dataset to confirm that our results are independent of datasets to some extent. The alternative dataset was downloaded from GEO (GSE16441). The procedures applied to analyze the GEO dataset are similar to those applied to the dataset obtained from TCGA. The only difference is the number of samples, miRNAs, and mRNAs. After repeating the same procedures, we realized that 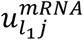 and 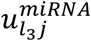 (*l*_1_ = *l*_3_ *= 2*) also varied between normal and tumor samples (Fig 1). *P*-values computed by the t-test were 6.74 × 10^−22^ for mRNA and 2.54 × 10^−18^ for miRNA. In order to ascertain whether 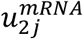 and 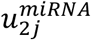 are significantly correlated, we calculated the *PCC* between them, which was 0. 931 (p-value = 1.58 × 10^−15^), indicating that they are highly correlated.

Next, we checked if the selected miRNAs and mRNAs were common between the TCGA and GEO datasets. We identified three miRNAs – hsa-miR-141, hsa-miR-210, and hsa-miR-200c, which are listed in Table 1. On the other hand, 209 genes were identified. After restricting genes included in both TCGA and GEO datasets, we evaluated the overlap as the confusion matrix (Table 2).

**Table 2.** Confusion matrix between genes selected in TCGA and GEO dataset.

|  |  | GEO |  |
| --- | --- | --- | --- |
|  |  | Not selected | Selected |
| TCGA | Not selected | 17209 | 160 |
|  | Selected | 60 | 11 |

The *P*-value determined using the Fisher exact test was 8.97 × 10^−11^ and the odds ratio was 19.7. Therefore, the coincidence between selected genes in the TCGA and GEO datasets is significant, and the results obtained for TCGA are robust and not highly dependent upon specific samples.

To test the superiority over the conventional methods, we applied the t-test, SAM [44], and limma [45] to the TCGA and GEO datasets, respectively. After applying these statistical methods, *P*-values were calculated and adjusted based on the BH criterion. Then, 13,895 genes and 399 miRNAs for TCGA and 12,152 genes and 78 miRNAs for GEO were associated with adjusted *P*-values less than 0.01 by *t*-test. At the same time, by SAM, 14,485 genes and 441 miRNAs for TCGA and 16,336 genes and 108 miRNAs for GEO were selected. Finally, limma selected 18,225 genes and 662 miRNAs for TCGA and 28,524 genes and 319 miRNAs for GEO. Relative to the TD method, the t-test, SAM, and limma identified a larger number of genes and miRNAs using the *P*-values as criteria. If the top ranked (small enough or restricted) number of genes and miRNAs was selected by the t-test, SAM and limma, the coincidence between TCGA and GEO might be compatible. Therefore, we selected the same number of genes and miRNAs by the t-test, SAM and limma as those selected by TD. Only one miRNA and no genes were common between the TCGA and GEO datasets for *t*-test. We could not reduce the number of genes and miRNAs selected by SAM, since it attributed *P* = 0 to more genes and miRNAs than those selected by TD for both TCGA and GEO, as did limma for miRNA. Limma could select a reduced number of genes for TCGA and GEO, while no common genes were selected between them. Therefore, we determined that the t-test, SAM, and limma could identify less coincident sets of genes and miRNAs between TCGA and GEO. In conclusion, this strongly suggests that the proposed method is superior to the t-test, SAM, and limma.

Although we cannot deny the possibility that more advanced or sophisticated methods can compete with TD-based unsupervised FE, since this method was fast, simple, and robust and gave us a biologically reasonable set of genes and miRNAs, we believe that the present results are worth publishing, even without more comprehensive comparisons with methods other than limma, SAM, and t-test.

We did not apply PCA to miRNAs and mRNAs separately and instead applied TD to a tensor that was generated by merging these two. To demonstrate this point, we applied PCA to miRNAs and mRNAs retrieved from TCGA. We noticed that the second PC loading attributed to miRNA and mRNA samples, 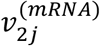 and 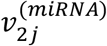, are not only mutually correlated but also distinct between tumours and normal tissues; the *PCC* between them, which was 0.839 (*P*-value = 2.74 × 10^−87^), is less significant than that when TD was employed (0.905, P = 1.63 × 10^−121^). *T*-test applied to 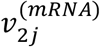 and 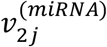 to evaluate significant distinctions between tumours and normal tissues gave us *P* = 2.33 × 10^−36^ for mRNA and *P* = 2.39 × 10^−77^ for miRNA, which are at best comparable ( *P* = 7.10 × 10^−39^ for mRNA and *P* = 2.13 × 10^−71^ for miRNA when TD was employed). To evaluate if PCA could select common genes and miRNAs between TCGA and GEO, we also applied PCA to the GEO dataset and selected 10 miRNAs and 70 mRNAs for TCGA and three miRNAs and 131 mRNAs for GEO. Since three miRNAs selected for GEO were also included in 10 miRNA selected for TCGA, coincidence between GEO and TCGA is the same for miRNAs between TD and PCA. Conversely, since we could find only six genes in common between TCGA and GEO, coincidence for mRNAs is less than with TD, which identified 11 genes in common. In conclusion, although miRNAs and mRNAs can also be successfully identified separately by PCA, the integrated analysis of TD has some advantages over PCA.

## Conclusions

In this study, we applied the TD-based unsupervised FE method to the KIRC miRNA expression and gene expression data. The TD-based method can identify miRNA signatures with differential expression between normal tissues and tumors as well as significant correlations between the gene expression data. Selected mRNAs and miRNAs are not only mutually correlated but are also significantly related to various aspects of cancers. This suggests that integrated analysis performed by TD-based unsupervised FE is an effective strategy; it can identify biologically significant pairs of miRNAs and mRNAs despite its simplicity, which is not easy by other strategies.

## Materials and Methods

### Tensors and tensor decomposition (TD)

Tensor [17] is a mathematical structure for storing datasets associated with more than two properties. If we measure miRNA and mRNA expression for the samples, we cannot avoid storing these two measurements into two separate matrices. However, by using tensor we can store these two datasets into a tensor, because tensors can have more than two suffixes, which matrices do not have.

TD [17] is a mathematical trick that can approximate tensors as the summation of series whose terms are expressed via the outer product of vectors, each of which represent individual property (in this specific example, these vectors correspond to mRNAs, miRNAs, and samples).

### Tensor decomposition methods

Figure 7 shows how TD and PCA were applied to miRNA and mRNA expression to select critical miRNAs and mRNAs for KIRC. The miRNAseq and mRNAseq expression data for KIRC were retrieved from the TCGA Data Portal Research Network (https://gdcportal.nci.nih.gov/).

**Fig. 7.**
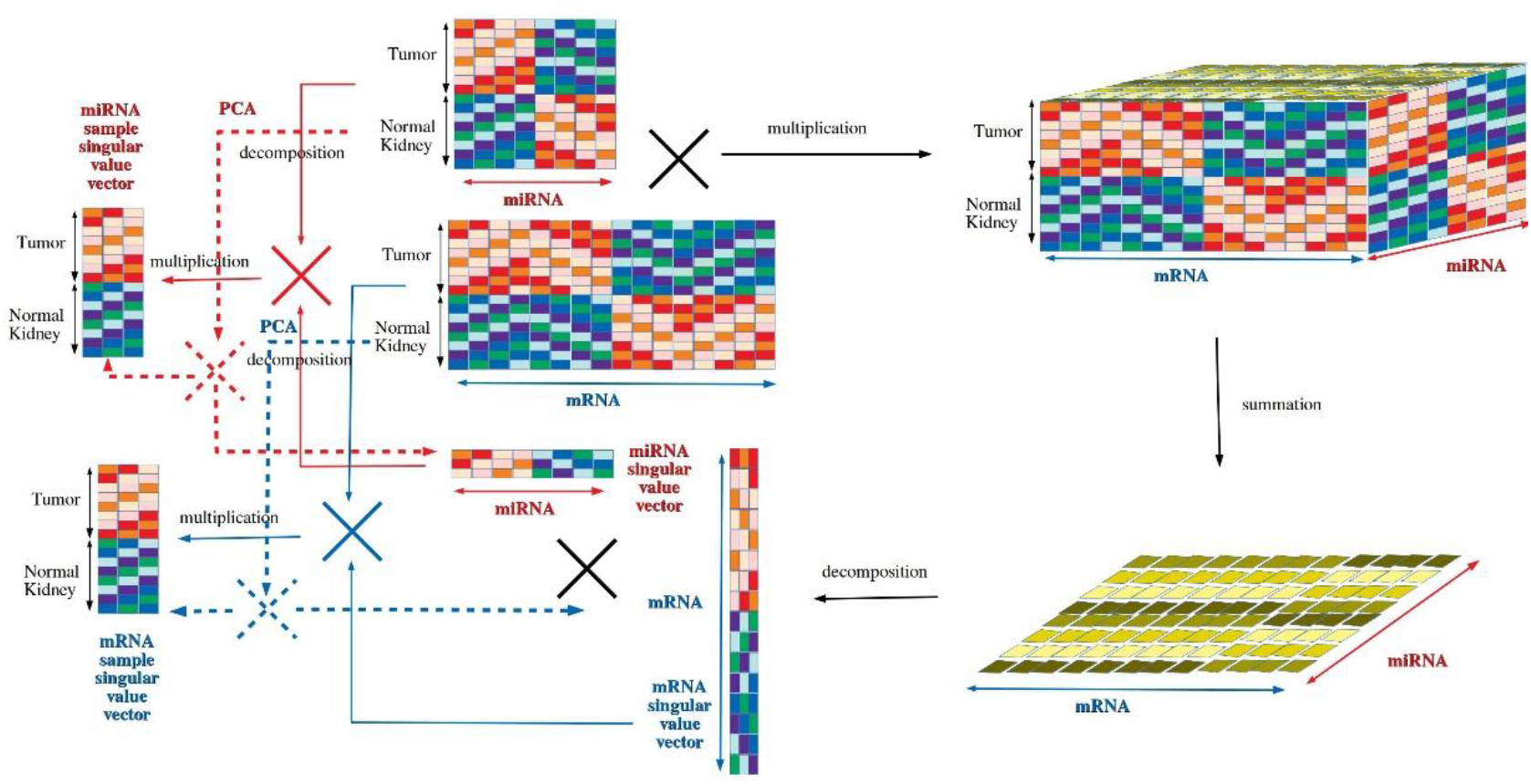
Illustration of TD and PCA application to mRNAs and miRNAs expression, respectively. TD workflow (solid line): mRNA and miRNA expression provided as matrices (middle upper) were multiplied to generate a tensor (right upper), which was converted to a matrix with summation sample suffix (lower right). The tensor was decomposed into a product of miRNA and mRNA singular value matrices (middle lower), which was converted to miRNA sample and mRNA sample singular value vectors (left). PCA workflow (broken lines): PCA directly decomposed miRNA and mRNA expression matrices into a product of PC loading attributed to samples and PC scores attributed to miRNA or mRNA.

TD is a natural extension of matrix factorization and is regarded as a generalization of the singular value decomposition (SVD) method. It is a useful technique uncovering the underlying low-dimensional structures in the tensor. There are two popular tensor decomposition algorithms: canonical polyadic decomposition (CPD) and Tucker decomposition [46]. The rank decomposition method, CPD, is to express a tensor as the sum of a finite number of rank-one tensors. The Tucker decomposition decomposes a tensor into a so-called core tensor and multiple matrices.

TD-based unsupervised FE was applied to analyze mRNA and miRNA expression profiles. Let *xij*^*(mRNA)*^ denote the expression profiles of the *i*th mRNA (*i* = 1, …*N*) of the *jth* sample ( *j* = 1, … *M*), whereas *xkj*^*(miRNA)*^ denotes the expression profiles of the *k*th miRNA ( *k* = 1, …*K*) of the *j*th sample ( *j* = 1, … *M*). Next, we generated a tensor, that is,

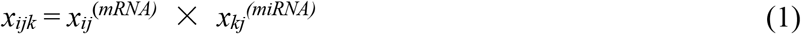

*xijk* is subjected to Tucker decomposition as follows:

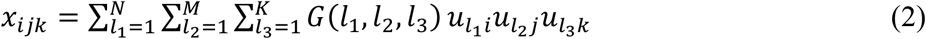

where G ∈ *R*^*N*×*M*×*K*^ is the core tensor and *u*_*l*_1_*i*_ ∈ *R*^*N*×*N*^, *u*_*l*_2_*j*_ ∈ *R*^*M*×*M*^ and *u*_*l*_ *u*_*l*_3_*k*_ ∈ *R*^*K*×*K*^ are singular value matrices that are orthogonal. Because Tucker decomposition is not unique, we have to specify how Tucker decomposition was derived. In particular, we chose higher-order singular value decomposition (HOSVD). Given that *xijk* ∈ *R*^*N*×*M*×*K*^ is too large (*N* × *M* × *K* = 19536 × 324 × 825 ≅ 5.22 × 10^9^; for actual numbers of *N*, *M*, and *K*, see below) to apply TD, we generated a matrix, which is given by:

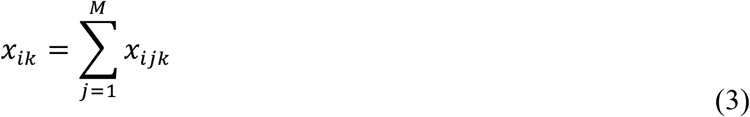

By applying SVD, we can get *u*_*l*_1_*j*_ and *u*_*l*_3_*k*_ as

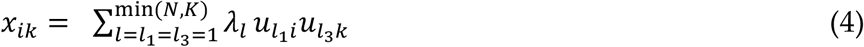

Then, we can also obtain two *u*_*l*_2_*k*_ that correspond to miRNA and mRNA expression:

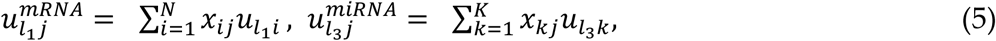

Selection of genes can be determined using the following quantities,

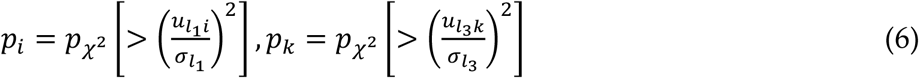

where p_ϰ^2^_ [>x] is the cumulative probability that the argument is greater than *x* in a ϰ^2^ distribution. *σ*_*l*_1__ and *σ*_*l*_1__ denote the standard deviations for *u*_*l*_3_i_ and *u*_*l*_3_k_, respectively. After the P-values are adjusted by means of the Benjamini–Hochberg (BH) criterion, miRNAs and mRNAs that are associated with adjusted P-values less than 0.01 are selected as those showing differences in expression between controls (normal tissues) and treated samples (tumors).

Although the reason CPD was not employed is fully described in my recent book **[17]**, I briefly describe it here. First of all, CPD cannot give us unique solutions, only initial value-dependent solutions that prevent us from interpreting the results uniquely. Another disadvantage of CPD compared with HOSVD is computation time: CPD is ten times slower than HOSVD [19]. Thus, there is no reason to employ CPD over HOSVD.

Although many other algorithms can compute Tucker decomposition, to our knowledge, HOSVD is the fastest method. In addition, since it works well applied to various problems [17] as well as in the present study, there is no need to employ other algorithms besides HOSVD.

### Principal component analysis methods

Similar to TD, PCA can also be applied to miRNA and miRNA expression, although separately rather than in an integrated manner. *xij*^(*mRNA)*^ and *xkj*^(*miRNA)*^ are normalized such that 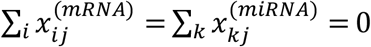 and 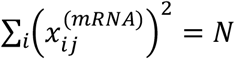, 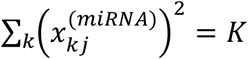. The *l*th PC score 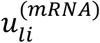, attributed to *i*th mRNA, can be obtained as the eigenvector of the gram matrix, 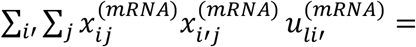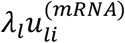, where *λ*_*l*_ is the eigenvalue. The *l*th PC loading, 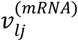, attributed to the *j*th sample, can be obtained by 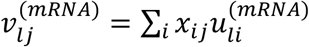. The exact same procedure was applied to 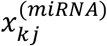, resulting in 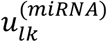 and 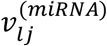. After identifying that and 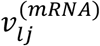 were 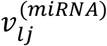 mutually correlated as well as distinct between tumour and normal tissues, corresponding 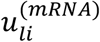 and 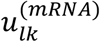 were used to attribute *P*-values to *i*th mRNAs and *k*th miRNAs, respectively, with distribution as was done with TD. Finally, mRNAs and miRNAs associated with adjusted *P*-values of less than 0.01 were selected.

### mRNA and miRNA expression

Expression profiles of the mRNA and miRNA were retrieved from the Firebrowse database (http://firebrowse.org/). The samples consisted of 253 kidney tumors and 71 normal kidney tissues (*M* = 324). The number of mRNAs measured was *N* = 19536, and the number of measured miRNAs was *K* = 825. Another dataset was downloaded from the GEO database (https://www.ncbi.nlm.nih.gov/geo/) with GEO ID GSE16441, and two files, GSE16441-GPL6480_series_matrix.txt.gz (for mRNA) and SE16441-GPL8659_series_matrix.txt.gz (for miRNA), were used. A total of *N* = 33698 mRNAs and *K* = 319 miRNAs were measured for 17 patients and 17 healthy controls (*M* = 34).

### Analysis of the correlation between miRNA and gene expression

Correlations between 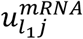 and 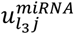 (*l*_1_ = *l*_3_= 2) as well as 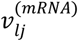 and 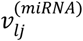 (*l*=2) were quantified by the Pearson’s correlation coefficient (*PCC*). The *PCC* and P-values were calculated using the *corr* function and *cor.test* function in the R software, respectively.

### Biological function analysis

We evaluated the biological significance of the set of differentially expressed miRNAs and their correlated mRNAs. Biological annotations of the prognostic miRNAs and mRNAs were examined by employing the DIANA-miRPath [47] and MSigDB [48] databases, respectively.

## Supporting information

Supplementary plots (Kaplan Meier plots)

## Supplementary Materials

Supplementary figures. The results of the Kaplan-Meier plots of the 23 KIRC survival-associated genes by using OncoLnc [31]. Supplementary Tables S1 to S4 that include numerical data as well as detailed descriptions that correspond to Figs. 3 to 6.

## Acknowledgments

Dr. Ka-Lok Ng is funded by the Ministry of Science and Technology, Taiwan (MOST), grant number MOST 108-2221-E-468-020, and also supported by the Asia University, grant numbers 105-asia-11, 106-asia-06, 106-asia-09, 107-asia-02 and 107-asia-09. Dr. Y-h Taguchi is supported by Kakenhi 20H04848, 20K12067, 19H05270 and 17K00417. We would like to thank Editage (http://www.editage.com) for English language editing.

## Author Contributions

Ka-Lok Ng foresee the research, prepared the data, writing—original draft preparation, review and editing. Y-h Taguchi performed the formal analysis, writing—original draft preparation, review and editing.

## Availability of Materials and Data

All the raw data were publicly available, which were obtained from the Firebrowse database (http://firebrowse.org/) and the GEO database (https://www.ncbi.nlm.nih.gov/geo/).

## References

1. Vargas-Rondon, N., Villegas, V. E. & Rondon-Lagos, M. The role of chromosomal instability in cancer and therapeutic responses. Cancers (Basel). 10, 4 (2017).

2. Hanahan, D. & Weinberg, R. A. Hallmarks of cancer: the next generation. Cell. 144, 646–74 (2011).

3. Feinberg, A.P. & Vogelstein, B. Hypomethylation distinguishes genes of some human cancers from their normal counterparts. Nature. 301, 89–92 (1983).

4. Rouhi, A., Mager, D. L., Humphries, R. K. & Kuchenbauer, F. MiRNAs, epigenetics, and cancer. Mamm. Genome. 19, 517–25 (2008).

5. Medina, P. P. & Slack, F. S. microRNAs and cancer: an overview. Cell Cycle. 7, 2485–2492 (2008).

6. Zhang, W., Dahlberg, J. E. & Tam, W. MicroRNAs in tumorigenesis: a primer. Am. J. Pathol. 171, 728–38 (2007).

7. Inamoto, T. et al. A panel of microRNA signature as a tool for predicting survival of patients with urothelial carcinoma of the bladder. Dis. Markers. 2018, 5468672 (2018).

8. Shi, X. H. et al. A five-microRNA signature for survival prognosis in pancreatic adenocarcinoma based on TCGA data. Sci. Rep. 8, 7638 (2018).

9. Yu, Y., Feng, X. & Cang, S. A two-microRNA signature as a diagnostic and prognostic marker of pancreatic adenocarcinoma. Cancer Manag. Res.. 10, 1507–1515 (2018).

10. Zhou, H. et al. A panel of eight-miRNA signature as a potential biomarker for predicting survival in bladder cancer. J. Exp. Clin. Cancer Res. 34, 53 (2015).

11. Lokeshwar, S. D. et al. Molecular characterization of renal cell carcinoma: a potential three-microRNA prognostic signature. Cancer Epidemiol. Biomarkers Prev. 27, 464–472 (2018).

12. Luo, Y., Chen, L., Wang, G., Xiao, Y., Ju, L. & Wang, X. Identification of a three-miRNA signature as a novel potential prognostic biomarker in patients with clear cell renal cell carcinoma. J. Cell. Biochem. 120, 13751–13764 (2019).

13. Xie, M. et al. Identification and validation of a four-miRNA (miRNA-21-5p, miRNA-9-5p, miR-149-5p, and miRNA-30b-5p) prognosis signature in clear cell renal cell carcinoma. Cancer Manag. Res. 10, 5759–5766 (2018).

14. Taguchi, Y.-h. One-class differential expression analysis using tensor decomposition-based unsupervised feature extraction applied to integrated analysis of multiple omics data from 26 lung adenocarcinoma cell lines. 2017 IEEE 17th International Conference on Bioinformatics and Bioengineering (BIBE). 131–138 (2017).

15. Taguchi, Y.-h. & Ng, K. *[Regular Paper]* Tensor decomposition–based unsupervised feature extraction for integrated analysis of TCGA data on microRNA expression and promoter methylation of genes in ovarian cancer. 2018 IEEE 18th International Conference on Bioinformatics and Bioengineering (BIBE). 195–200 (2018).

16. Taguchi, Y.-h. Tensor decomposition based unsupervised feature extraction applied to bioinformatics *in Application of omics, AI and blockchain in bioinformatics research* (eds. Tsai, J.-P. & Ng, K.-L.) (World Scientifc Publisher, 2019).

17. Taguchi, Y.-h. Unsupervised feature extraction applied to bioinformatics: A PCA based and TD based approach. Springer International (2019).

18. Taguchi, Y.-h. Multiomics data analysis using tensor decomposition based unsupervised feature extraction –comparison with DIABLO– in *Intelligent computing theories and application*. ICIC 2019. Lecture notes in computer science vol. 11643. (eds. Huang D.-S., Bevilacqua, V. & Premaratne, P.) (Springer, 2019).

19. Taguchi, Y.-h. & Turki, T. Neurological disorder drug discovery from gene expression with tensor decomposition. Curr. Pharm. Des. 25,4589–4599.

20. Taguchi, Y.-h. Tensor decomposition-based unsupervised feature extraction identifies candidate genes that induce post-traumatic stress disorder-mediated heart diseases. BMC Med. Genomics. 10, s67 (2017).

21. Taguchi, Y.-h. Identification of candidate drugs using tensor-decomposition-based unsupervised feature extraction in integrated analysis of gene expression between diseases and DrugMatrix datasets. Sci. Rep. 7, 13733 (2017).

22. Taguchi, Y.-h. Tensor decomposition-based unsupervised feature extraction applied to matrix products for multi-view data processing. PLoS One. 12, e0183933 (2017).

23. Taguchi, Y.-h. Tensor decomposition-based unsupervised feature extraction can identify the universal bature of sequence-bonspecific off-target regulation of mRNA mediated by microRNA transfection. Cells. 7, 54 (2018).

24. Taguchi, Y.-h. Tensor decomposition-based and principal-component-analysis-based unsupervised feature extraction applied to the gene expression and methylation profiles in the brains of social insects with multiple castes. BMC Bioinformatics. 19, 99 (2018).

25. Taguchi, Y.-h. Correction: Tensor decomposition-based unsupervised feature extraction applied to matrix products for multi-view data processing. PLoS One. 13, e0200451 (2018).

26. Taguchi, Y.-h. Drug candidate identification based on gene expression of treated cells using tensor decomposition-based unsupervised feature extraction for large-scale data. BMC Bioinformatics. 19, 388 (2019).

27. Taguchi, Y.-h. & Turki, T. Tensor decomposition-based unsupervised feature extraction applied to single-cell gene expression analysis. Front Genet. 10, 864 (2019).

28. Torres, V. E. & Harris, P. C. Strategies targeting cAMP signaling in the treatment of polycystic kidney disease. J. Am. Soc. Nephrol. 25, 18–32 (2014).

29. Piazzon, N., Maisonneuve, C., Guilleret, I., Rotman, S., Constam, D. B. Bicc1 links the regulation of cAMP signaling in polycystic kidneys to microRNA-induced gene silencing. J. Mol. Cell Biol. 4, 398–408 (2012).

30. Isakoff, M. S. et al. Inactivation of the Snf5 tumor suppressor stimulates cell cycle progression and cooperates with p53 loss in oncogenic transformation. Proc. Natl. Acad. Sci. U S A. 102, 17745–17750 (2005).

31. Sarnowska, E. et al. Evaluation of the role of downregulation of SNF5/INI1 core subunit of SWI/SNF complex in clear cell renal cell carcinoma development. Am. J. Cancer Res. 7, 2275–2289 (2017).

32. Zang, Y. et al. Eukaryotic translation initiation factor 3b is both a promising prognostic biomarker and a potential therapeutic target for patients with clear cell renal cell carcinoma. J. Cancer. 8, 3049–3061 (2017).

33. Jaiswal, P. K., Koul, S., Palanisamy, N. & Koul, H. K. Eukaryotic translation initiation factor 4 gamma 1 (eIF4G1): a target for cancer therapeutic intervention? Cancer Cell Int. 19, 224 (2019).

34. Solarek, W., Koper, M., Lewicki, S., Szczylik, C. & Czarnecka, A. M. Insulin and insulin-like growth factors act as renal cell cancer intratumoral regulators. J. Cell. Commun. Signal. 13, 381–394 (2019).

35. Braczkowski, R., Białożyt, M., Plato, M., Mazurek, U. & Braczkowska, B. Expression of insulin-like growth factor family genes in clear cell renal cell carcinoma. Contemp. Oncol. (Pozn). 20, 130–136 (2016).

36. Tracz, A. F., Szczylik, C., Porta, C., Czarnecka, A. M. Insulin-like growth factor-1 signaling in renal cell carcinoma. BMC Cancer. 16, 453 (2016).

37. Major, J. M., Pollak, M. N., Snyder, K., Virtamo, J. & Albanes, D. Insulin-like growth factors and risk of kidney cancer in men. Br. J. Cancer. 103, 132–135 (2010).

38. Lee, M. & Rhee, I. Cytokine cignaling in tumor progression. Immune Netw. 17, 214–227 (2017).

39. Doehn, C., Kausch, I., Melz, S., Behm, A. & Jocham, D. Cytokine and vaccine therapy of kidney cancer. Expert Rev. Anticancer Ther. 4, 1097–1111 (2004).

40. Macleod, L. C. et al. Trends in metastatic kidney cancer survival from the cytokine to the targeted therapy era. Urology. 86, 262–268 (2015).

41. Unver, N. & McAllister, F. IL-6 family cytokines: Key inflammatory mediators as biomarkers and potential therapeutic targets. Cytokine Growth Factor Rev. 41, 10–17 (2018).

42. Kaminska, K., Czarnecka, A. M., Escudier, B., Lian, F. & Szczylik, C. Interleukin-6 as an emerging regulator of renal cell cancer. Urol. Oncol. 33, 476–485 (2015).

43. Ishibashi, K. et al. Interleukin-6 induces drug resistance in renal cell carcinoma. Fukushima J. Med. Sci. 64, 103–110 (2018).

44. Tusher, V. G., Tibshirani, R. & Chu, G. Significance analysis of microarrays applied to the ionizing radiation response. Proc. Natl. Acad. Sci. U S A. 98, 5116–5121 (2001).

45. Ritchie, M. E. et al. Limma powers differential expression analyses for RNA-sequencing and microarray studies. Nucleic Acids Res. 43, e47–e47 (2015).

46. Rabanser, S., Shchur, O. & Günnemann, S. Introduction to tensor decompositions and their applications in machine learning. Preprint at https://arxiv.org/pdf/1711.10781.pdf (2017).

47. Vlachos, I. S. et al. DIANA-miRPath v3.0: deciphering microRNA function with experimental support. Nucleic Acids Res. 43, W460–W466 (2015).

48. Liberzon, A., Birger, C., Thorvaldsdóttir, H., Ghandi, M., Mesirov, J. P., Tamayo, P. The Molecular Signatures Database (MSigDB) hallmark gene set collection. Cell Syst. 1, 417–425 (2015).

