## Supplementary Tables for "Identification of miRNA signatures for kidney renal clear cell carcinoma using the tensor-decomposition method"

**Table S1.** The top 10 oncogenic signatures of the 72 genes reported by the MSigDB. #Genes (K): the number of genes in each overexpressed gene set. # Genes in overlap (*k*): overlaps with genes selected via the TD-based unsupervised FE method. .

| **Gene Set Name**  **[# Genes (K)]** | **Description** | **#Genes in overlap (*k*)** | **p-value** | **FDR q-value** |
| --- | --- | --- | --- | --- |
| [CAMP_UP.V1_UP](http://software.broadinstitute.org/gsea/msigdb/geneset_page.jsp?geneSetName=CAMP_UP.V1_UP)  [[200](http://software.broadinstitute.org/gsea/msigdb/download_geneset.jsp?geneSetName=CAMP_UP.V1_UP&fileType=grp)] | Genes up-regulated in primary thyrocyte cultures in response to cAMP signaling pathway activation by thyrotropin (TSH). | 7 | 9.97 e-8 | 1.88 e-5 |
| [SNF5_DN.V1_DN](http://software.broadinstitute.org/gsea/msigdb/geneset_page.jsp?geneSetName=SNF5_DN.V1_DN)  [[168](http://software.broadinstitute.org/gsea/msigdb/download_geneset.jsp?geneSetName=SNF5_DN.V1_DN&fileType=grp)] | Genes down-regulated in MEF cells (embryonic fibroblasts) with knockout of SNF5 [Gene ID=6598] gene. | 6 | 7.64 e-7 | 7.22 e-5 |
| ESC_V6.5_UP_  LATE.V1_UP  [188] | Genes up-regulated during the late stages of differentiation of embryoid bodies from V6.5 embryonic stem cells. | 6 | 1.47 e-6 | 9.27 e-5 |
| ESC_V6.5_UP_  EARLY.V1_DN  [[175](http://software.broadinstitute.org/gsea/msigdb/download_geneset.jsp?geneSetName=ESC_V6.5_UP_EARLY.V1_DN&fileType=grp)] | Genes down-regulated during the early stages of differentiation of embryoid bodies from V6.5 embryonic stem cells. | 5 | 1.98 e-5 | 8.54 e-4 |
| ESC_J1_UP_  LATE.V1_UP  [189] | Genes up-regulated during the late stages of differentiation of embryoid bodies from J1 embryonic stem cells. | 5 | 2.86 e-5 | 8.54 e-4 |
| [SIRNA_EIF4GI_UP](http://software.broadinstitute.org/gsea/msigdb/geneset_page.jsp?geneSetName=SIRNA_EIF4GI_UP)  [[95](http://software.broadinstitute.org/gsea/msigdb/download_geneset.jsp?geneSetName=SIRNA_EIF4GI_UP&fileType=grp)] | Genes up-regulated in MCF10A cells vs knockdown of the EIF4G1 [Gene ID=1981] gene by RNAi. | 4 | 3.11 e-5 | 8.54 e-4 |
| [P53_DN.V1_DN](http://software.broadinstitute.org/gsea/msigdb/geneset_page.jsp?geneSetName=P53_DN.V1_DN)  [[193](http://software.broadinstitute.org/gsea/msigdb/download_geneset.jsp?geneSetName=P53_DN.V1_DN&fileType=grp)] | Genes down-regulated in the NCI-60 panel of cell lines with mutated TP53 [Gene ID=7157]. | 5 | 3.16 e-5 | 8.54 e-4 |
| [MEL18_DN.V1_UP](http://software.broadinstitute.org/gsea/msigdb/geneset_page.jsp?geneSetName=MEL18_DN.V1_UP)  [[141](http://software.broadinstitute.org/gsea/msigdb/download_geneset.jsp?geneSetName=MEL18_DN.V1_UP&fileType=grp)] | Genes up-regulated in DAOY cells (medulloblastoma) upon knockdown of PCGF2 [Gene ID=7703] gene by RNAi. | 4 | 1.45 e-4 | 3.42 e-3 |
| [LTE2_UP.V1_UP](http://software.broadinstitute.org/gsea/msigdb/geneset_page.jsp?geneSetName=LTE2_UP.V1_UP)  [[188](http://software.broadinstitute.org/gsea/msigdb/download_geneset.jsp?geneSetName=LTE2_UP.V1_UP&fileType=grp)] | Genes up-regulated in MCF-7 cells (breast cancer) positive for ESR1 [Gene ID=2099] MCF-7 cells (breast cancer) and long-term adapted for estrogen-independent growth. | 4 | 4.33 e-4 | 8.51 e-3 |
| [RPS14_DN.V1_UP](http://software.broadinstitute.org/gsea/msigdb/geneset_page.jsp?geneSetName=RPS14_DN.V1_UP)  [[190](http://software.broadinstitute.org/gsea/msigdb/download_geneset.jsp?geneSetName=RPS14_DN.V1_UP&fileType=grp)] | Genes up-regulated in CD34^+^ hematopoietic progenitor cells after knockdown of RPS14 [Gene ID=6208] by RNAi. | 4 | 4.5 e-4 | 8.51 e-3 |

**Table S2.** The top 10 oncogenic signatures of the 72 genes reported by the MSigDB. #Genes (K): the number of genes in each overexpressed gene set. # Genes in overlap (k): overlaps with genes selected by TD-based unsupervised FE method.

| **Gene Set Name**  **[# Genes (K)]** | **Description** | **# Genes in overlap (k)** | **p-value** | **FDR**  **q-value** |
| --- | --- | --- | --- | --- |
| [REACTOME_REGULATION_OF_INSULIN_LIKE_GR_GROWTH_FACTOR_IGF_TRANSPORT_AND_UPTAK TAKE_BY_INSULIN_LIKE_GROWTH_FACTOR_BIN BINDING_PROTEINS_IGFBPS](http://software.broadinstitute.org/gsea/msigdb/geneset_page.jsp?geneSetName=REACTOME_REGULATION_OF_INSULIN_LIKE_GROWTH_FACTOR_IGF_TRANSPORT_AND_UPTAKE_BY_INSULIN_LIKE_GROWTH_FACTOR_BINDING_PROTEINS_IGFBPS) [[124](http://software.broadinstitute.org/gsea/msigdb/download_geneset.jsp?geneSetName=REACTOME_REGULATION_OF_INSULIN_LIKE_GROWTH_FACTOR_IGF_TRANSPORT_AND_UPTAKE_BY_INSULIN_LIKE_GROWTH_FACTOR_BINDING_PROTEINS_IGFBPS&fileType=grp)] | Regulation of insulin-like growth factor (IGF) transport and uptake by insulin-like growth factor binding proteins (IGFBPs) | 12 | 9.03 e-18 | 1.35 e-14 |
| [REACTOME_CYTOKINE_SIGNALING_IN_IMMUNE_ NE_SYSTEM](http://software.broadinstitute.org/gsea/msigdb/geneset_page.jsp?geneSetName=REACTOME_CYTOKINE_SIGNALING_IN_IMMUNE_SYSTEM) [[856](http://software.broadinstitute.org/gsea/msigdb/download_geneset.jsp?geneSetName=REACTOME_CYTOKINE_SIGNALING_IN_IMMUNE_SYSTEM&fileType=grp)] | Cytokine signaling within the immune system | 18 | 1.85 e-14 | 1.39 e-11 |
| [REACTOME_RESPONSE_TO_ELEVATED_PLATELET LET_CYTOSOLIC_CA2PLUS](http://software.broadinstitute.org/gsea/msigdb/geneset_page.jsp?geneSetName=REACTOME_RESPONSE_TO_ELEVATED_PLATELET_CYTOSOLIC_CA2PLUS) [[132](http://software.broadinstitute.org/gsea/msigdb/download_geneset.jsp?geneSetName=REACTOME_RESPONSE_TO_ELEVATED_PLATELET_CYTOSOLIC_CA2PLUS&fileType=grp)] | Response to elevated platelet cytosolic Ca^2+^ | 9 | 3.42 e-12 | 1.71 e-9 |
| [REACTOME_SIGNALING_BY_INTERLEUKINS](http://software.broadinstitute.org/gsea/msigdb/geneset_page.jsp?geneSetName=REACTOME_SIGNALING_BY_INTERLEUKINS) [[631](http://software.broadinstitute.org/gsea/msigdb/download_geneset.jsp?geneSetName=REACTOME_SIGNALING_BY_INTERLEUKINS&fileType=grp)] | Signaling by interleukins | 13 | 1.53 e-10 | 4.86 e-8 |
| [REACTOME_INNATE_IMMUNE_SYSTEM](http://software.broadinstitute.org/gsea/msigdb/geneset_page.jsp?geneSetName=REACTOME_INNATE_IMMUNE_SYSTEM) [[1104](http://software.broadinstitute.org/gsea/msigdb/download_geneset.jsp?geneSetName=REACTOME_INNATE_IMMUNE_SYSTEM&fileType=grp)] | Innate immune system | 16 | 1.62 e-10 | 4.86 e-8 |
| [REACTOME_PLATELET_ACTIVATION_SIGNALING ING_AND_AGGREGATION](http://software.broadinstitute.org/gsea/msigdb/geneset_page.jsp?geneSetName=REACTOME_PLATELET_ACTIVATION_SIGNALING_AND_AGGREGATION) [[260](http://software.broadinstitute.org/gsea/msigdb/download_geneset.jsp?geneSetName=REACTOME_PLATELET_ACTIVATION_SIGNALING_AND_AGGREGATION&fileType=grp)] | Platelet activation, signaling and aggregation | 9 | 1.45 e-9 | 3.63 e-7 |
| [REACTOME_ENDOSOMAL_VACUOLAR_PATHWAY](http://software.broadinstitute.org/gsea/msigdb/geneset_page.jsp?geneSetName=REACTOME_ENDOSOMAL_VACUOLAR_PATHWAY) [[11](http://software.broadinstitute.org/gsea/msigdb/download_geneset.jsp?geneSetName=REACTOME_ENDOSOMAL_VACUOLAR_PATHWAY&fileType=grp)] | Endosomal/Vacuolar pathway | 4 | 3.63 e-9 | 7.78 e-7 |
| [REACTOME_GLUCONEOGENESIS](http://software.broadinstitute.org/gsea/msigdb/geneset_page.jsp?geneSetName=REACTOME_GLUCONEOGENESIS) [[34](http://software.broadinstitute.org/gsea/msigdb/download_geneset.jsp?geneSetName=REACTOME_GLUCONEOGENESIS&fileType=grp)] | gluconeogenesis | 5 | 5.22 e-9 | 9.79 e-7 |
| [REACTOME_POST_TRANSLATIONAL_PROTEIN_MO _MODIFICATION](http://software.broadinstitute.org/gsea/msigdb/geneset_page.jsp?geneSetName=REACTOME_POST_TRANSLATIONAL_PROTEIN_MODIFICATION) [[1429](http://software.broadinstitute.org/gsea/msigdb/download_geneset.jsp?geneSetName=REACTOME_POST_TRANSLATIONAL_PROTEIN_MODIFICATION&fileType=grp)] | Post-translational protein modification | 16 | 6.56 e-9 | 1.09 e-6 |
| [REACTOME_DISEASE](http://software.broadinstitute.org/gsea/msigdb/geneset_page.jsp?geneSetName=REACTOME_DISEASE) [[1075](http://software.broadinstitute.org/gsea/msigdb/download_geneset.jsp?geneSetName=REACTOME_DISEASE&fileType=grp)] | Disease | 14 | 1.02 e-8 | 1.53 e-6 |

**Table S3.** Survival analysis of KIRC using OncoLnc [31] (Kaplan plots are provided in the supplementary materials)

|  |  |  |  |  |  |  | Kaplan plot | | |
| --- | --- | --- | --- | --- | --- | --- | --- | --- | --- |
| Gene | Cox Coeff. | P-value | FDR Corrected | Rank | Median Expression | Mean Expression | Low (%) | High (%) | P-value |
| VWF | -0.3 | 1.90E-04 | 1.41E-03 | 2253 | 23278.72 | 25958.34 | 50 | 50 | 3.99E-03 |
| VEGFA | 0.25 | 2.90E-03 | 1.19E-02 | 4064 | 31629.77 | 35072.27 | 70 | 30 | 2.32E-02 |
| TMBIM6 | -0.2 | 5.00E-03 | 1.82E-02 | 4583 | 27241.35 | 28733.19 | 40 | 60 | 1.34E-02 |
| PODXL | -0.4 | 8.00E-06 | 1.22E-04 | 1092 | 6659.17 | 7271.06 | 50 | 50 | 1.36E-06 |
| PLVAP | -0.2 | 7.10E-03 | 2.39E-02 | 4946 | 15470.76 | 17515.66 | 50 | 50 | 4.10E-04 |
| PLIN2 | -0.3 | 6.20E-04 | 3.56E-03 | 2902 | 18947.56 | 22839.08 | 50 | 50 | 1.71E-05 |
| PCK1 | -0.3 | 1.00E-04 | 8.58E-04 | 1931 | 1120.74 | 3037.73 | 50 | 50 | 7.84E-06 |
| NDRG1 | -0.2 | 1.20E-02 | 3.61E-02 | 5506 | 50127.14 | 51689.99 | 60 | 40 | 2.72E-02 |
| ITM2B | -0.3 | 6.00E-04 | 3.47E-03 | 2880 | 34751.8 | 36807.63 | 50 | 50 | 1.36E-02 |
| HSPA8 | -0.3 | 1.20E-03 | 5.90E-03 | 3363 | 17668.96 | 18139.95 | 40 | 60 | 1.04E-02 |
| HLA-DRA | -0.2 | 3.80E-03 | 1.46E-02 | 4304 | 29068.65 | 32924.27 | 20 | 80 | 4.22E-02 |
| GATM | -0.3 | 4.20E-04 | 2.61E-03 | 2683 | 5433.14 | 6800.94 | 50 | 50 | 3.09E-04 |
| CYFIP2 | -0.5 | 2.20E-09 | 4.32E-07 | 82 | 3482.26 | 4051.73 | 50 | 50 | 9.88E-08 |
| CDH16 | -0.2 | 4.40E-03 | 1.65E-02 | 4430 | 4093.23 | 4940.33 | 50 | 50 | 1.14E-03 |
| CCND1 | -0.2 | 3.00E-03 | 1.22E-02 | 4068 | 17278.68 | 19256.81 | 50 | 50 | 2.85E-04 |
| ATP5B | -0.2 | 1.10E-02 | 3.37E-02 | 5360 | 11450.7 | 13211.83 | 30 | 70 | 2.59E-03 |
| ATP5A1 | -0.2 | 2.20E-03 | 9.54E-03 | 3812 | 7988.24 | 9278.65 | 50 | 50 | 2.86E-02 |
| ATP1B1 | -0.3 | 1.50E-03 | 7.03E-03 | 3514 | 18741.07 | 21002.32 | 50 | 50 | 3.90E-02 |
| ATP1A1 | -0.3 | 4.90E-05 | 4.98E-04 | 1634 | 12917.72 | 15392.31 | 40 | 60 | 2.34E-02 |
| AQP1 | -0.3 | 4.30E-05 | 4.52E-04 | 1580 | 16717.87 | 19036.22 | 50 | 50 | 3.11E-08 |
| APP | -0.4 | 1.90E-05 | 2.36E-04 | 1329 | 32137.14 | 33051.3 | 50 | 50 | 1.33E-06 |
| ALDOB | -0.3 | 4.40E-05 | 4.61E-04 | 1587 | 467.22 | 3374.03 | 50 | 50 | 3.27E-06 |
| AIF1L | -0.2 | 1.50E-03 | 7.03E-03 | 3510 | 1984.01 | 2798.36 | 60 | 40 | 2.80E-02 |

**Table S4.** The top 10 enriched KEGG pathways predicted by DIANA-mirpath for the 11 identified miRNAs (P-values are corrected). The full list can be obtained from http://snf-515788.vm.okeanos.grnet.gr/#mirnas=hsa-miR-210-3p;hsa-miR-210-5p;hsa-miR-891a-3p;hsa-miR-891a-5p;hsa-miR-200c-5p;hsa-miR-200c-5p;hsa-miR-141-5p;hsa-miR-141-3p;hsa-miR-122-3p;hsa-miR-122-5p;hsa-miR-155-3p;hsa-miR-155-5p;hsa-miR-508-3p;hsa-miR-508-5p;hsa-miR-514a-3p;hsa-miR-514a-5p;hsa-miR-184&methods=Tarbase;Tarbase;Tarbase;Tarbase;Tarbase;Tarbase;Tarbase;Tarbase;Tarbase;Tarbase;Tarbase;Tarbase;Tarbase;Tarbase;Tarbase;Tarbase;Tarbase&selection=0

| KEGG pathway | P-value | #genes | #miRNAs |
| --- | --- | --- | --- |
| Chronic myeloid leukemia | 5.90E-08 | 39 | 6 |
| Proteoglycans in cancer | 3.67E-06 | 72 | 8 |
| Prostate cancer | 2.58E-05 | 43 | 7 |
| Pathways in cancer | 3.10E-05 | 128 | 10 |
| Pancreatic cancer | 3.94E-05 | 32 | 5 |
| Glioma | 9.09E-05 | 28 | 5 |
| Hepatitis B | 9.11E-05 | 47 | 5 |
| Small cell lung cancer | 0.0002621 | 38 | 5 |
| Non-small cell lung cancer | 0.0002975 | 24 | 4 |
| Colorectal cancer | 0.0002975 | 28 | 7 |
| Endometrial cancer | 0.0007913 | 23 | 6 |
| Viral carcinogenesis | 0.0007913 | 59 | 8 |
| Bladder cancer | 0.001004 | 20 | 5 |
| Melanoma | 0.01584 | 25 | 5 |
| Renal cell carcinoma | 0.01613 | 27 | 5 |
| Hepatitis C | 0.02652153 | 44 | 6 |
