## Supplementary figures and images for "Identification of miRNA signatures for kidney renal clear cell carcinoma using the tensor-decomposition method"

### AIF1L_KIRC_83543_60_40.pdf

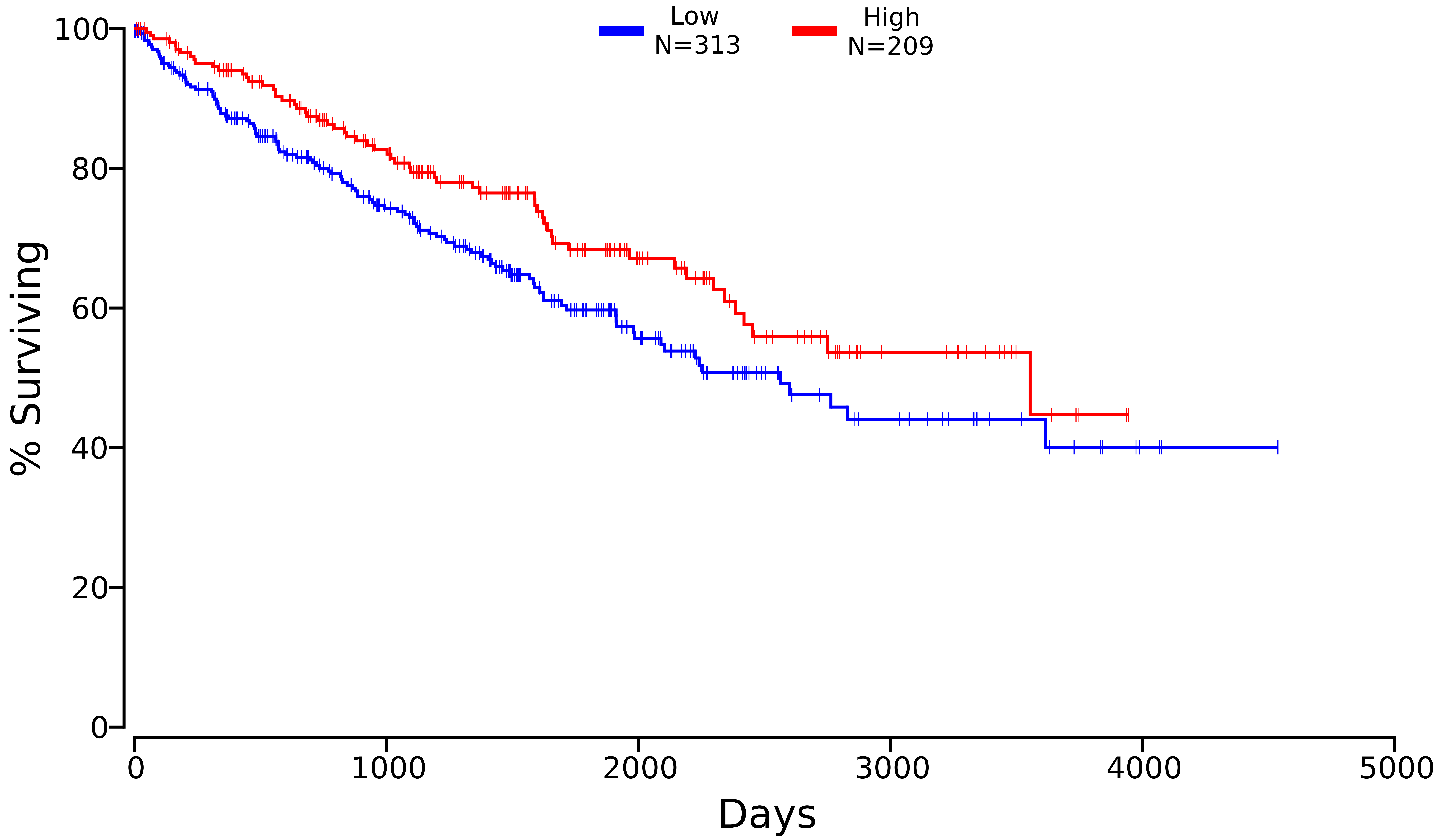

### ALDOB_KIRC_229_50_50.pdf

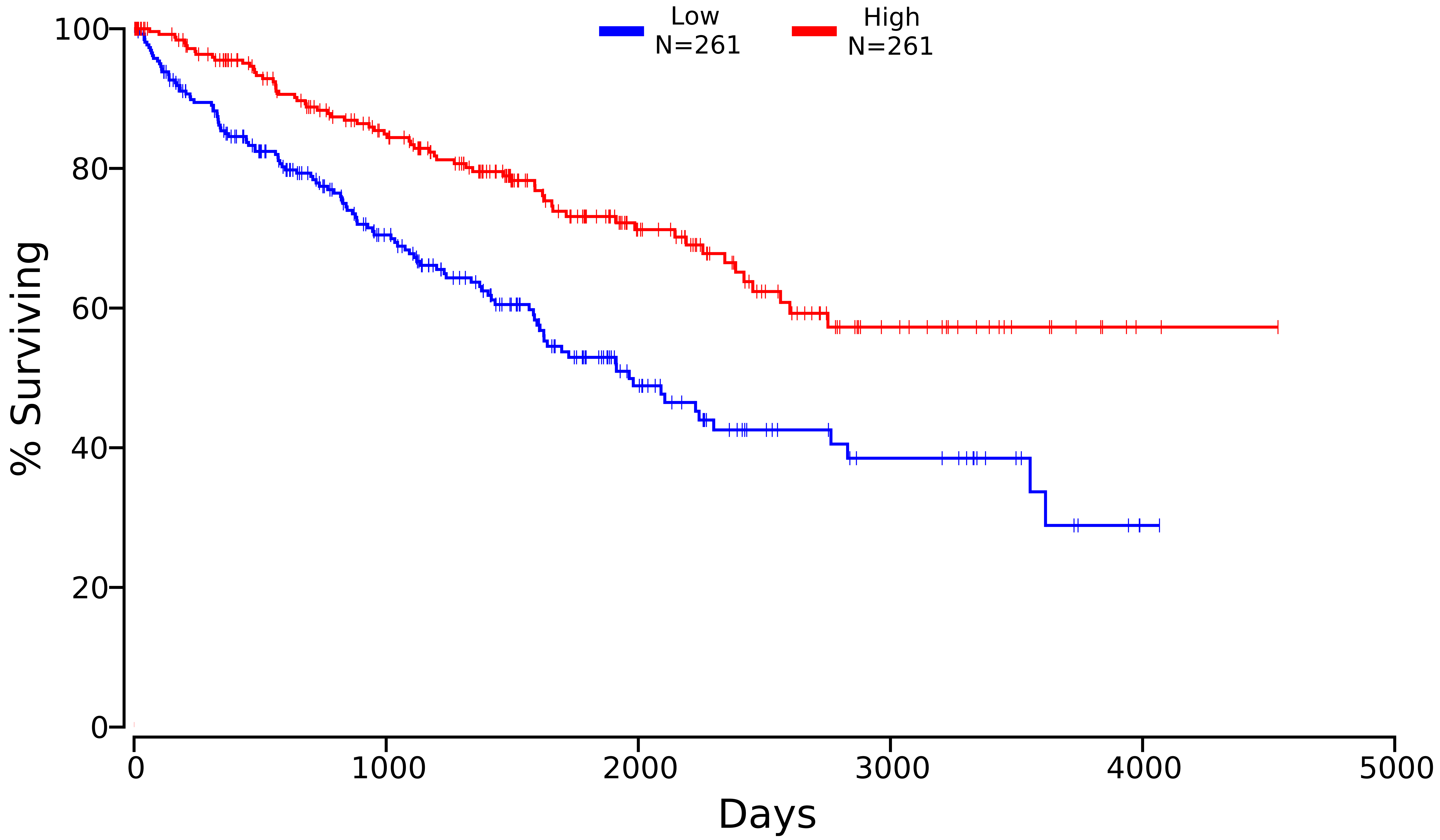

### APP_KIRC_351_50_50.pdf

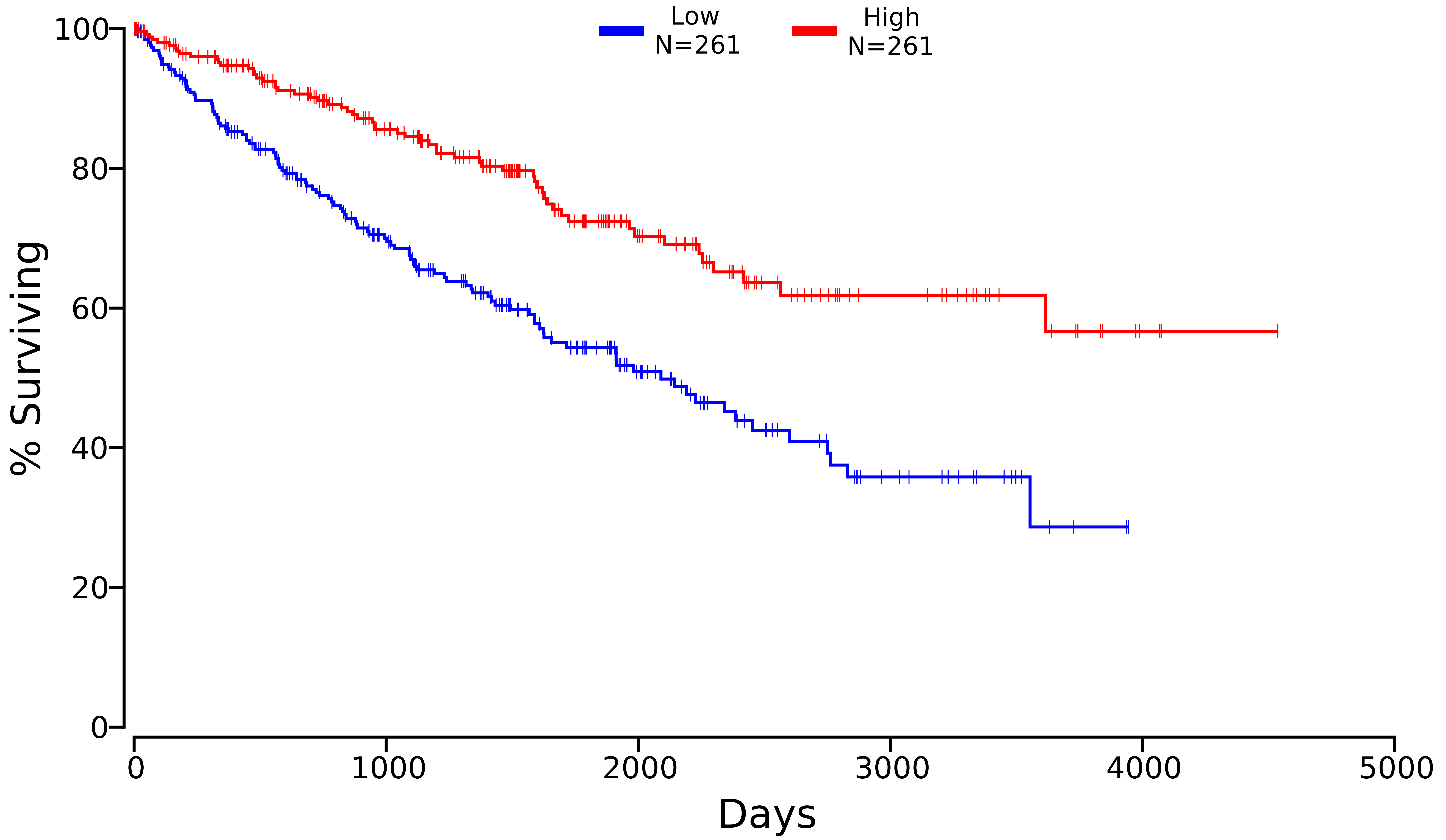

### AQP1_KIRC_358_50_50.pdf

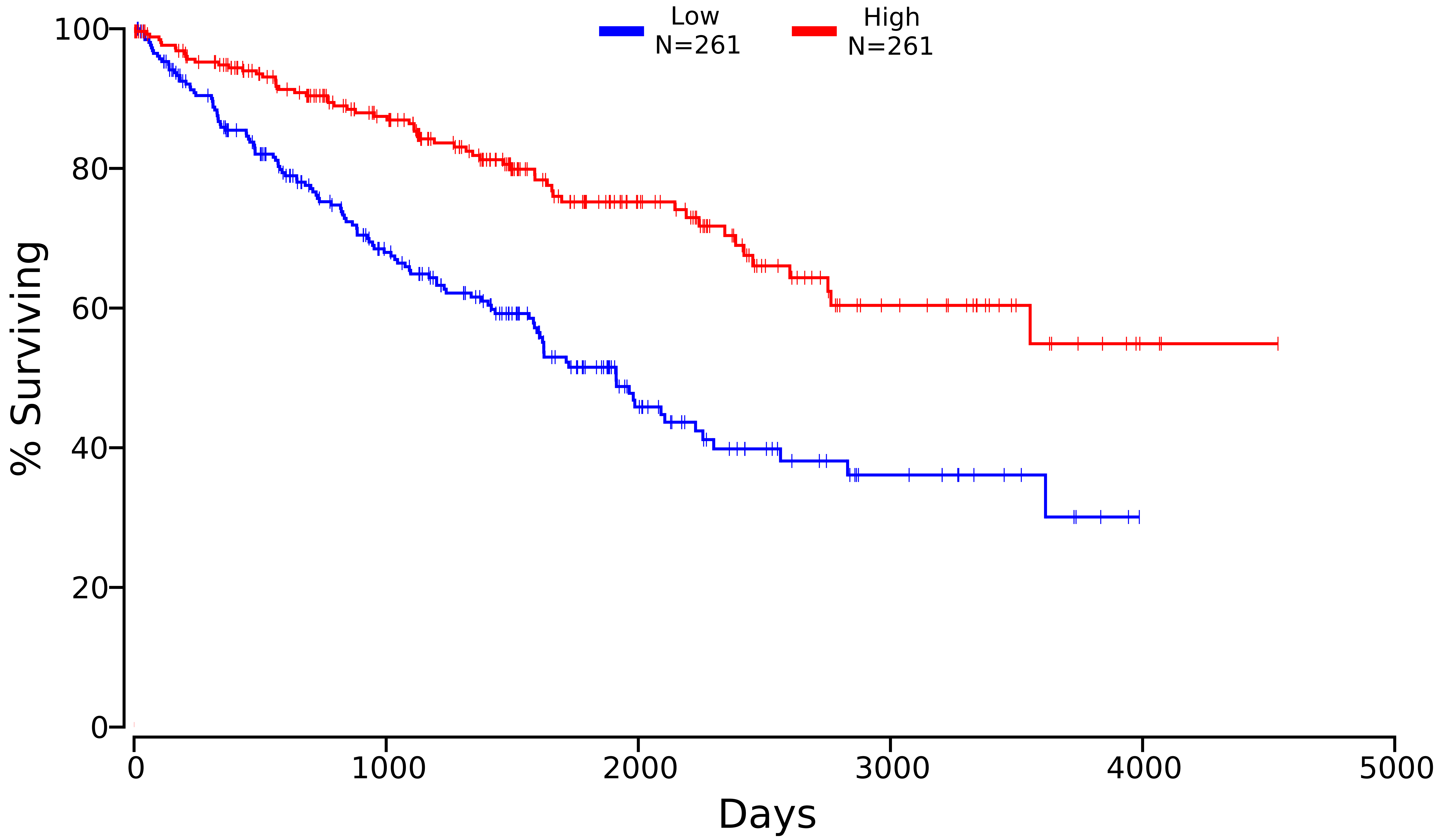

### ATP1A1_KIRC_476_40_60.pdf

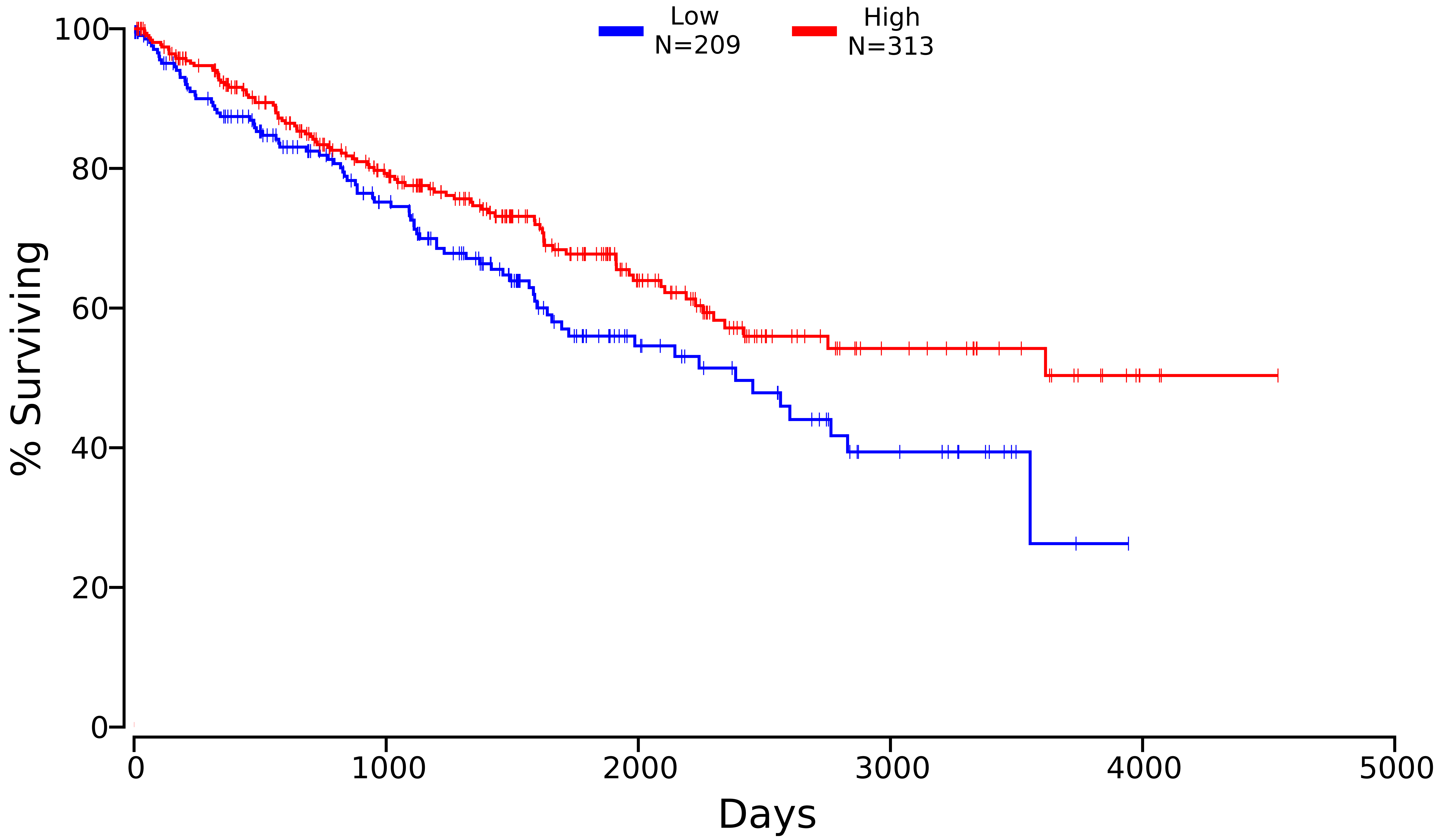

### ATP1B1_KIRC_481_50_50.pdf

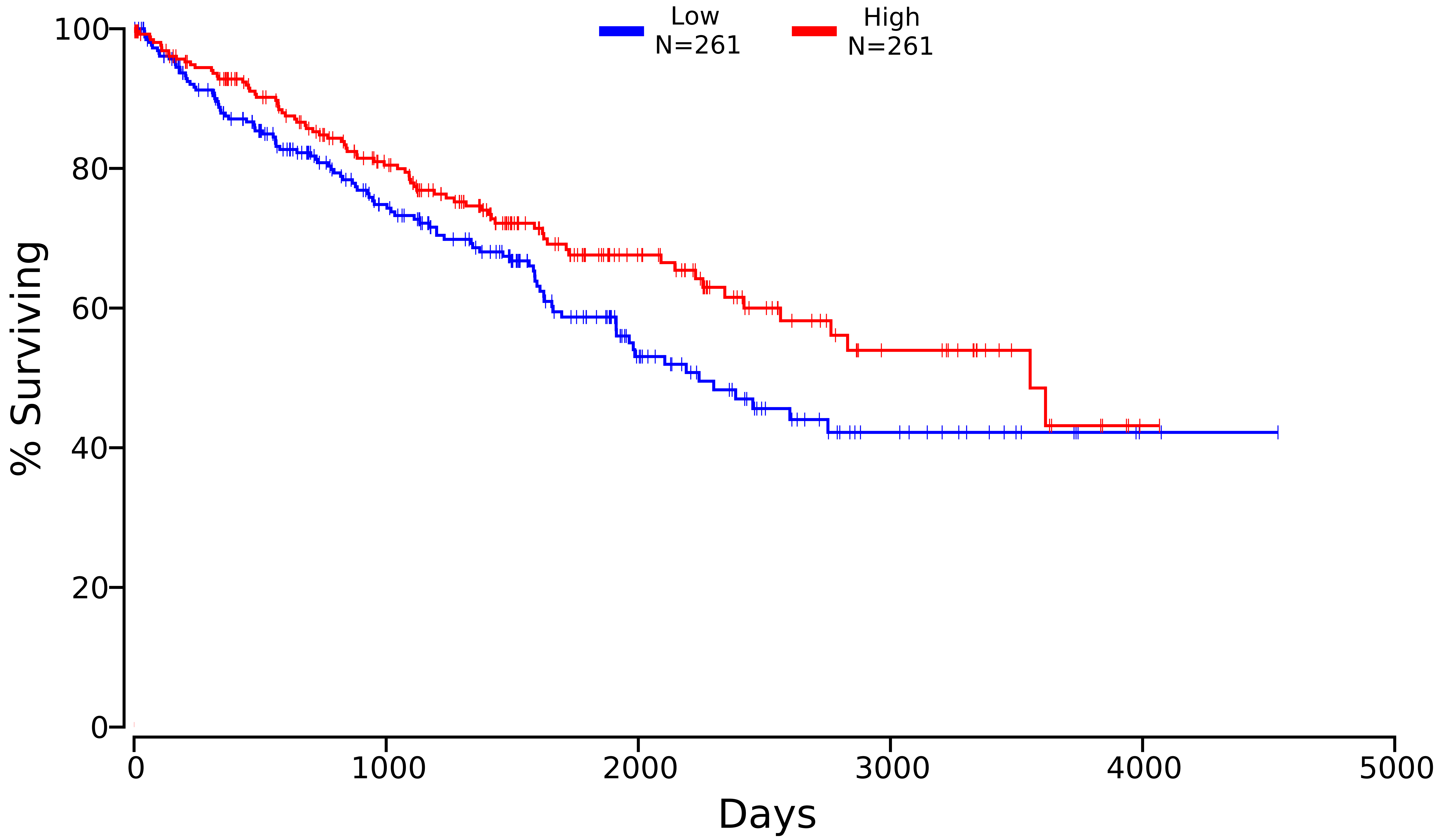

### ATP5A1_KIRC_498_50_50.pdf

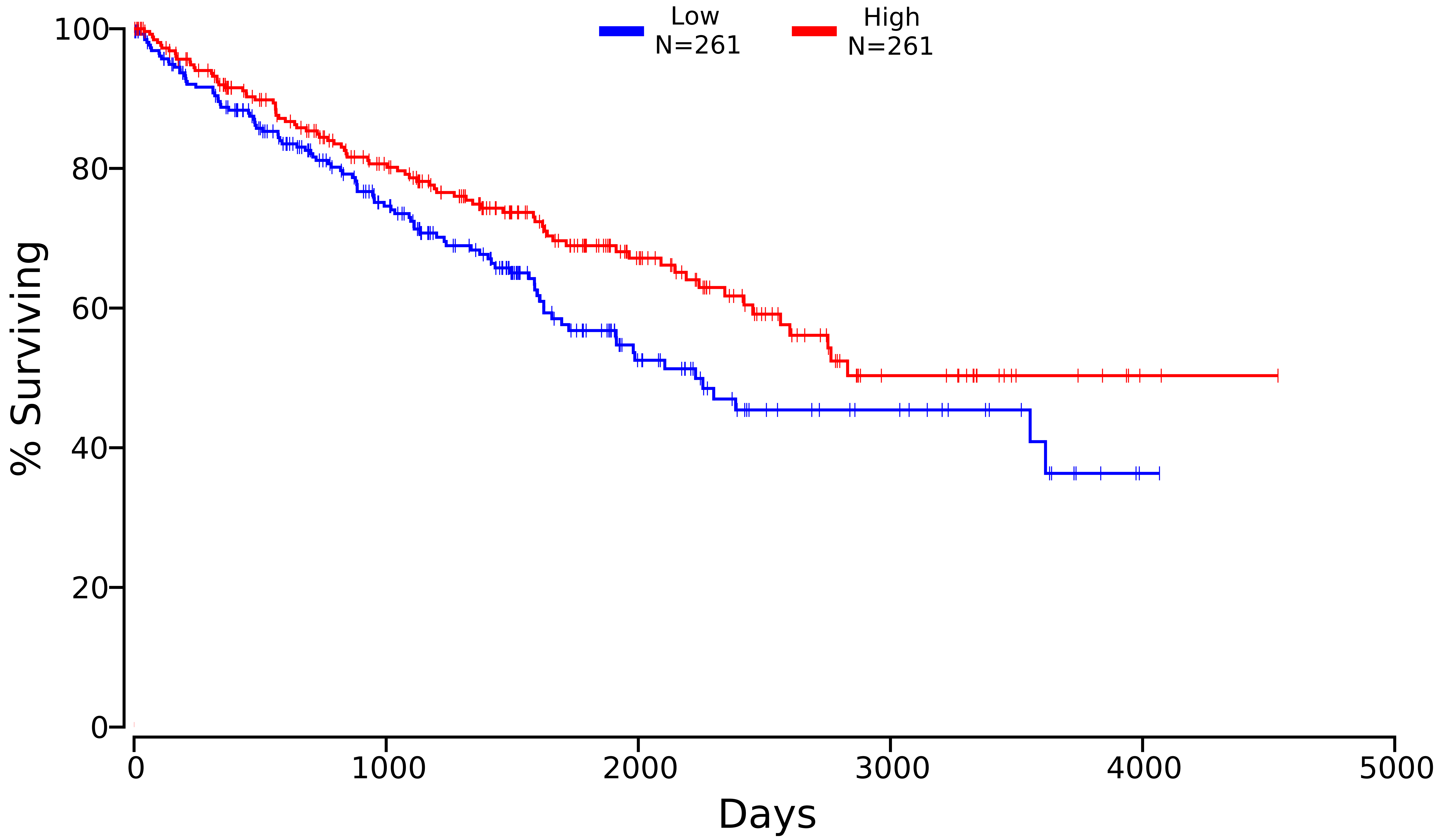

### ATP5B_KIRC_506_30_70.pdf

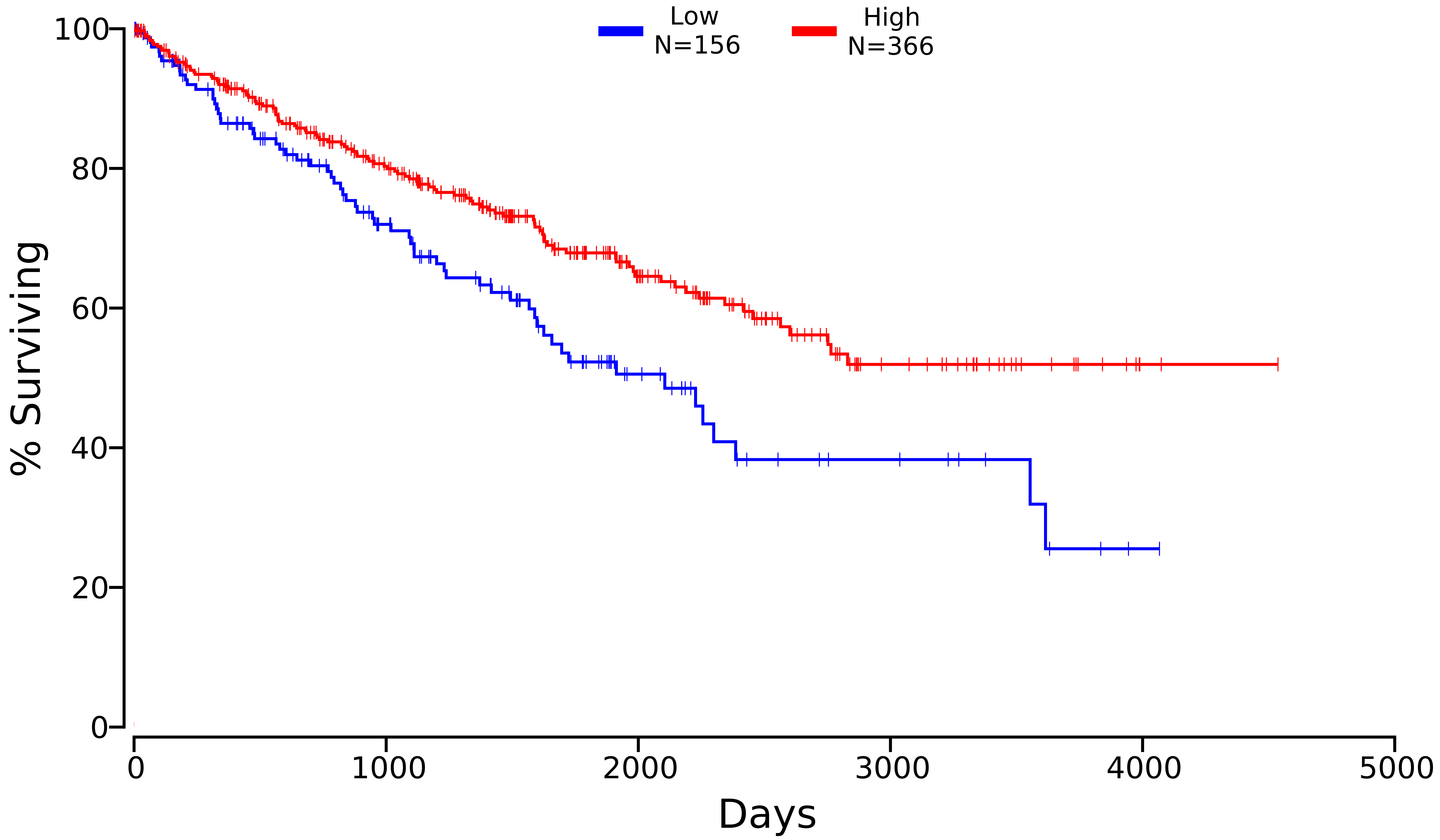

### CCND1_KIRC_595_50_50.pdf

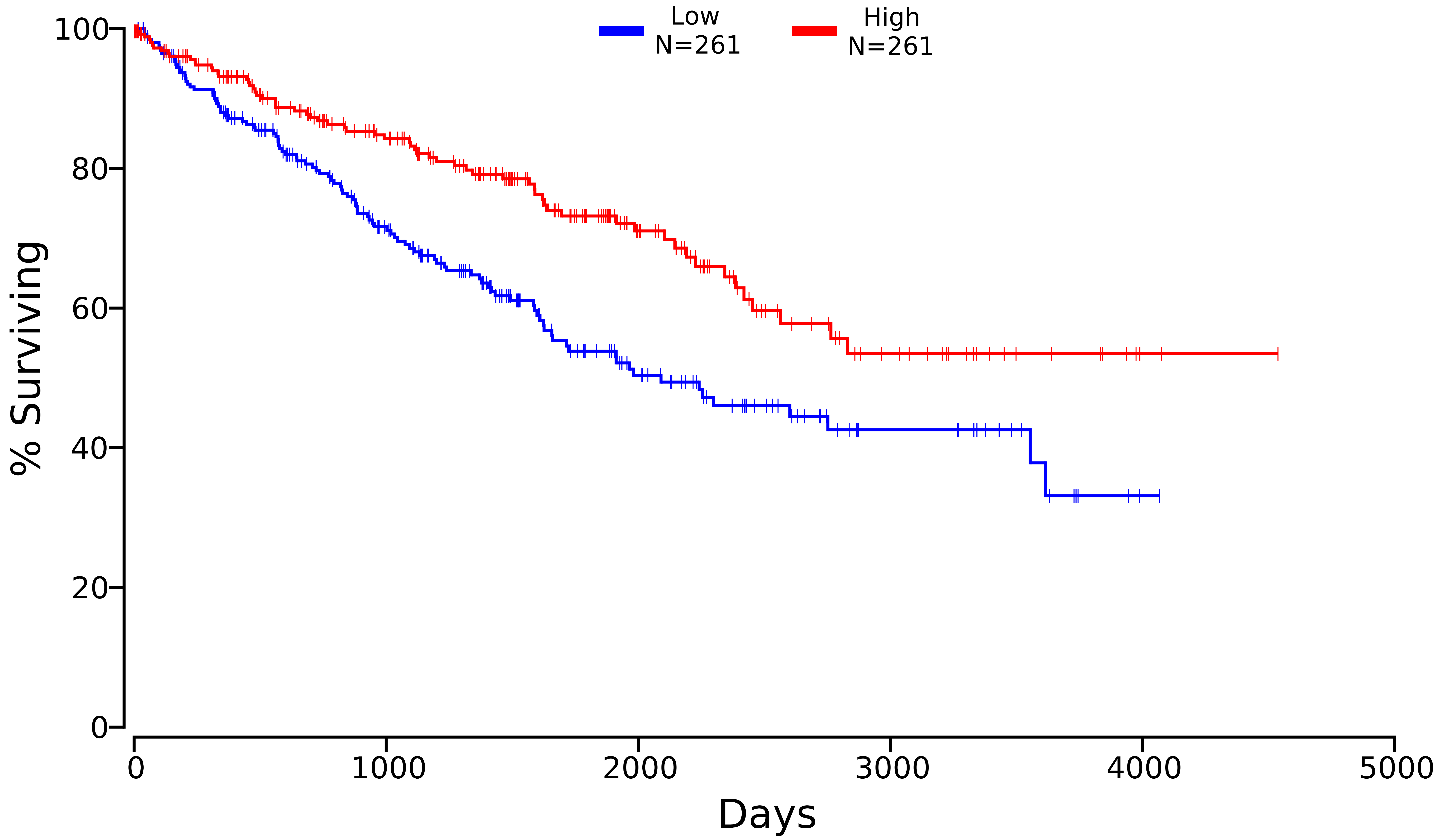

### CDH16_KIRC_1014_50_50.pdf

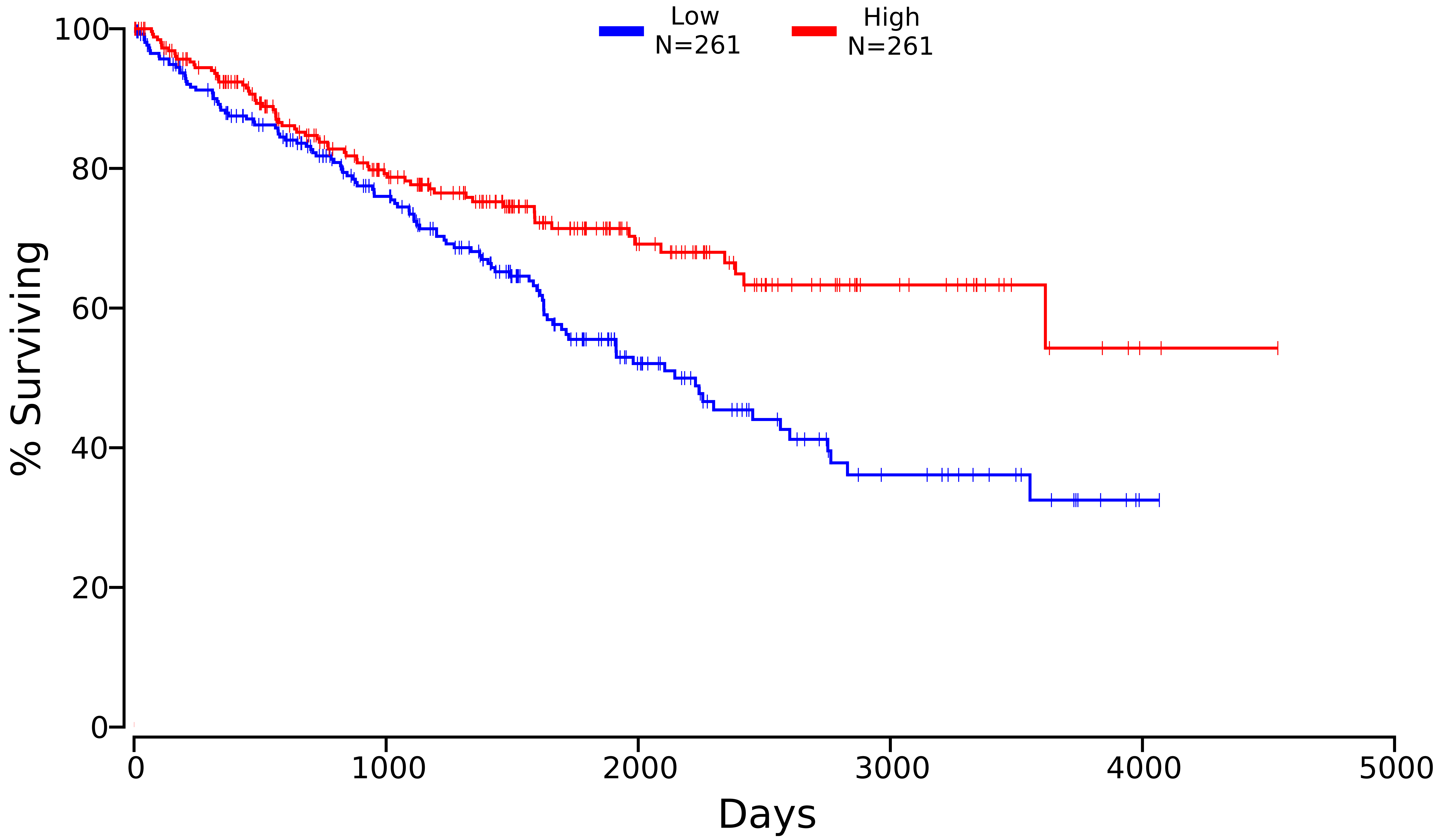

### CYFIP2_KIRC_26999_50_50.pdf

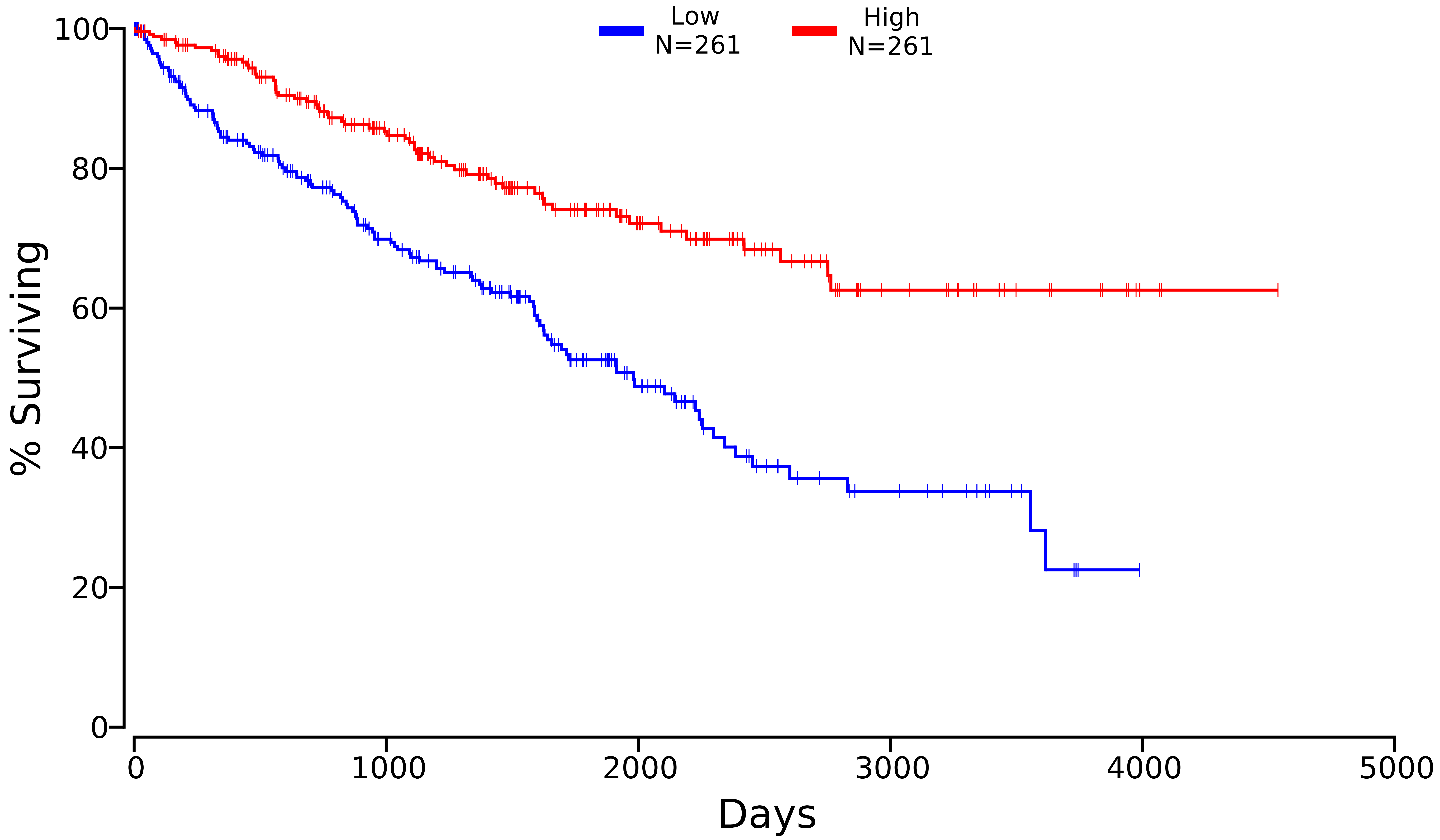

### GATM_KIRC_2628_50_50.pdf

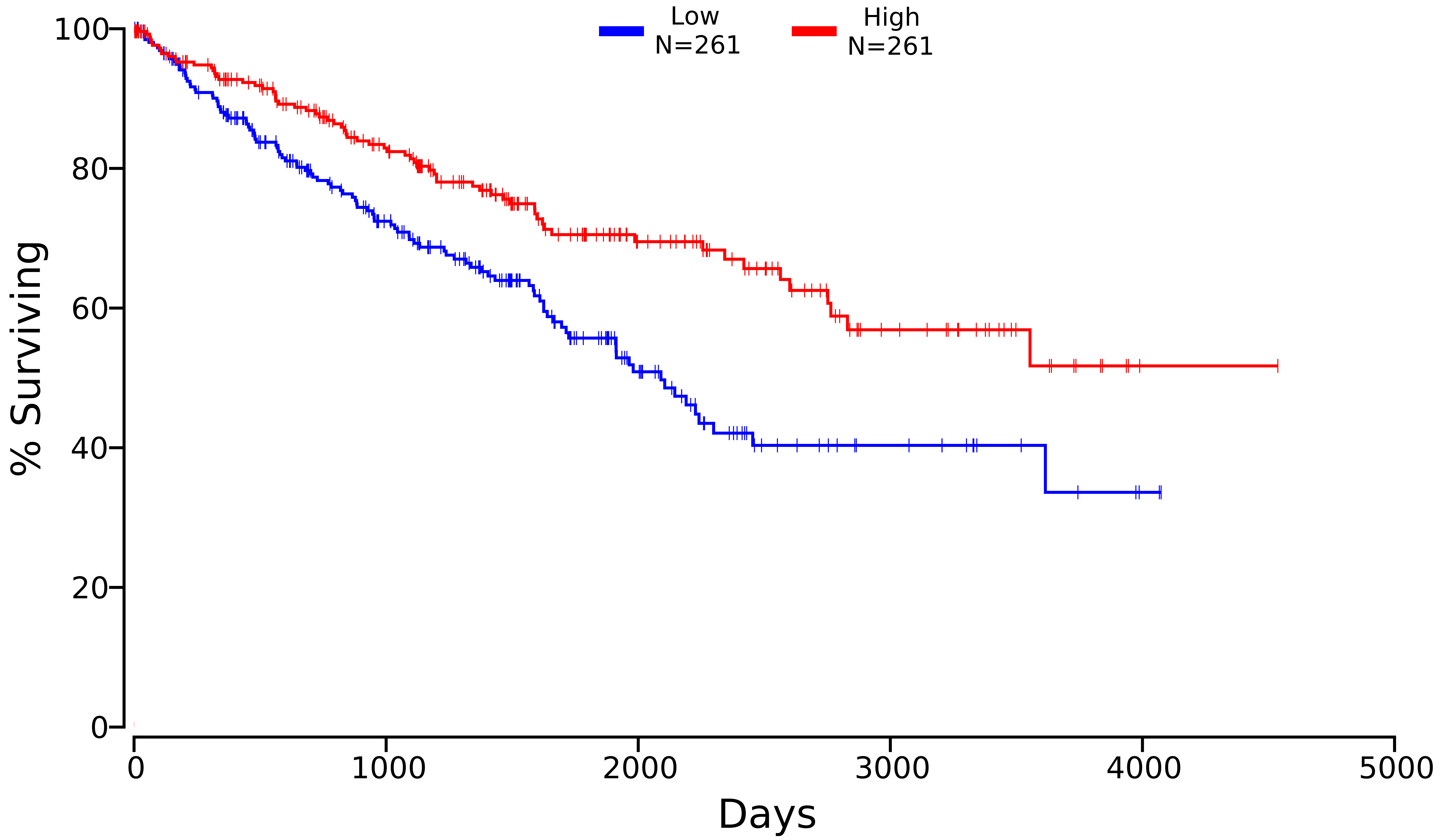

### HLA-DRA_KIRC_3122_20_80.pdf

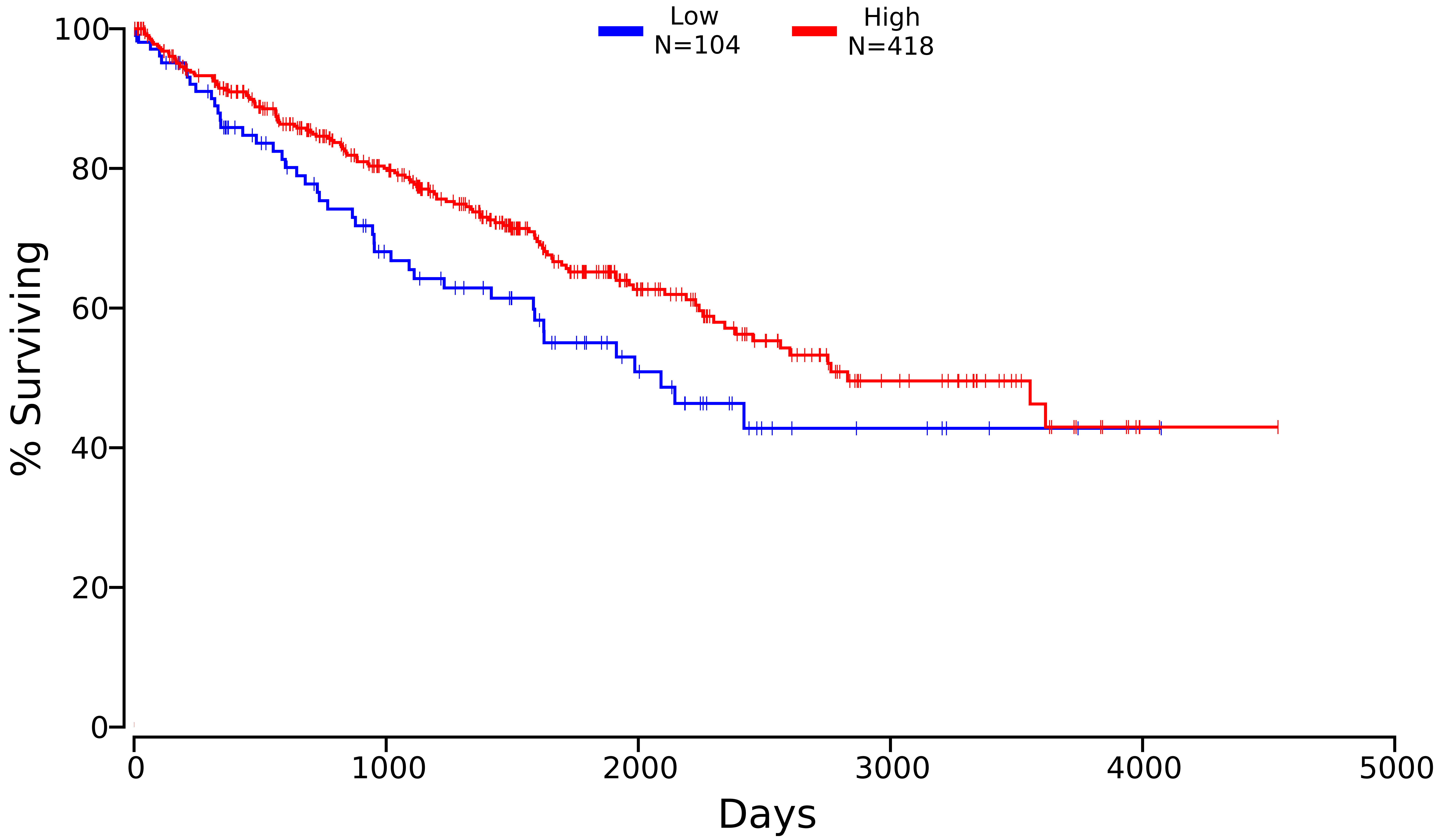

### HSPA8_KIRC_3312_40_60.pdf

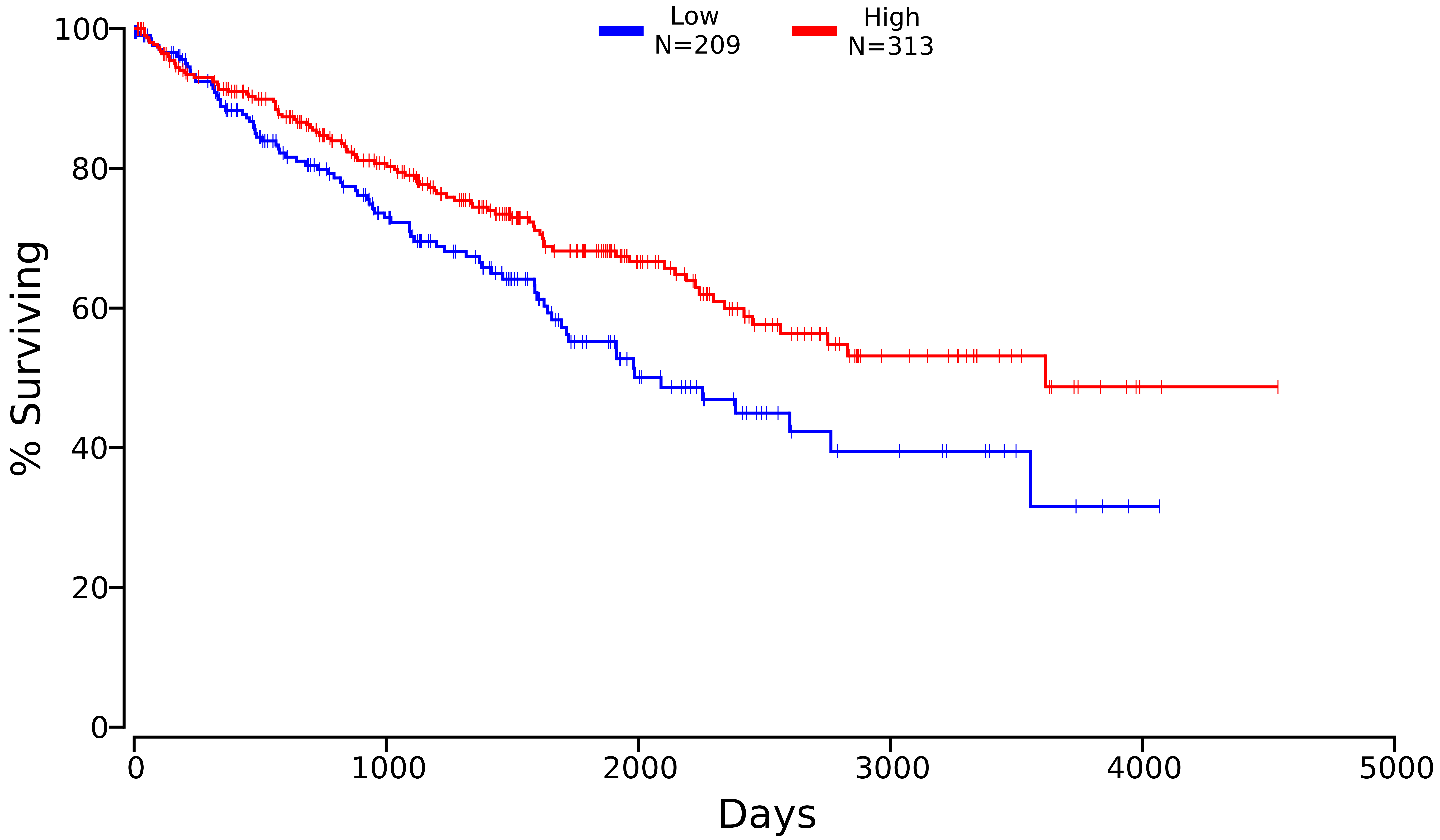

### ITM2B_KIRC_9445_50_50.pdf

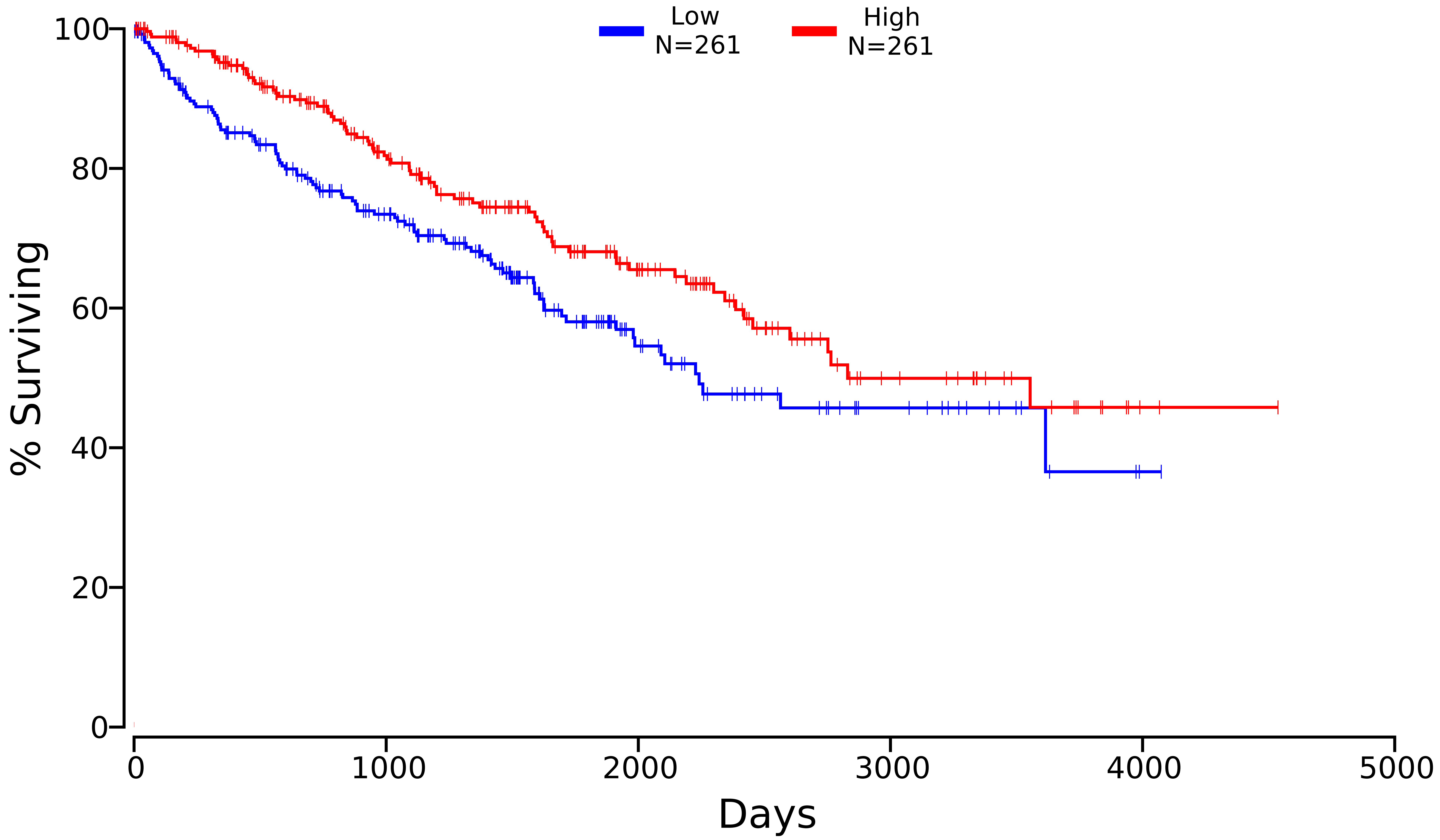

### NDRG1_KIRC_10397_60_40.pdf

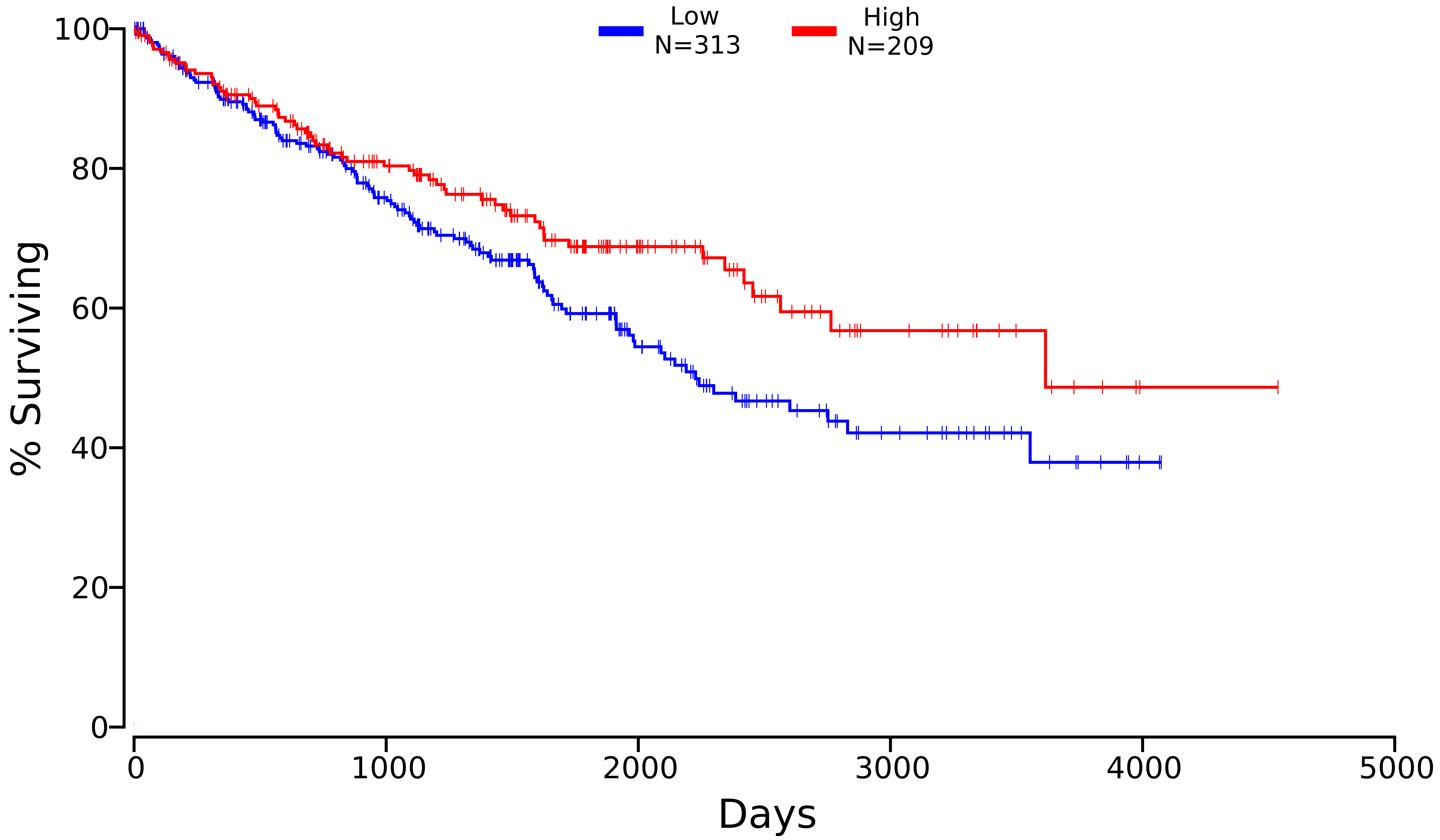

### PCK1_KIRC_5105_50_50.pdf

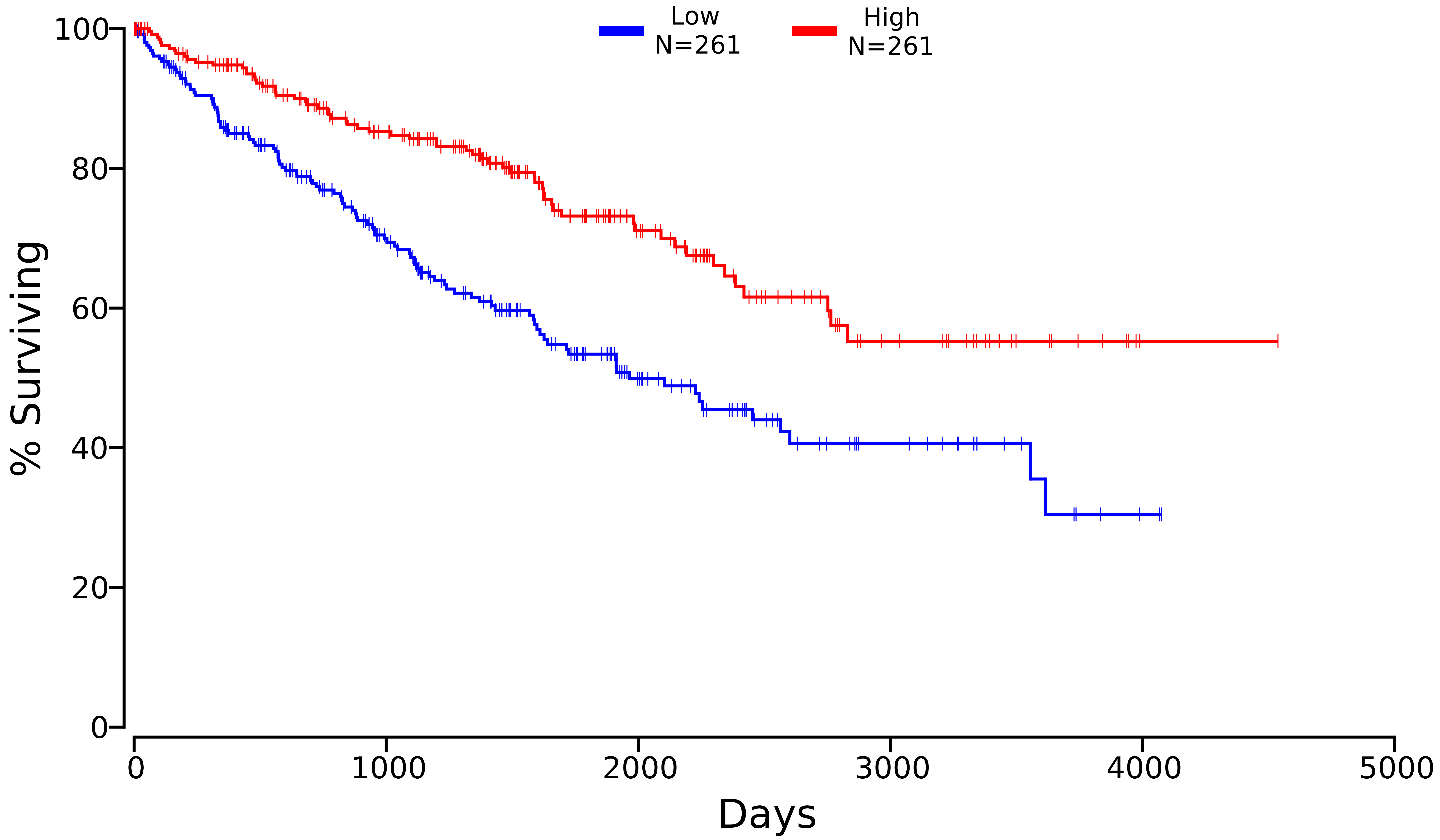

### PLIN2_KIRC_123_50_50.pdf

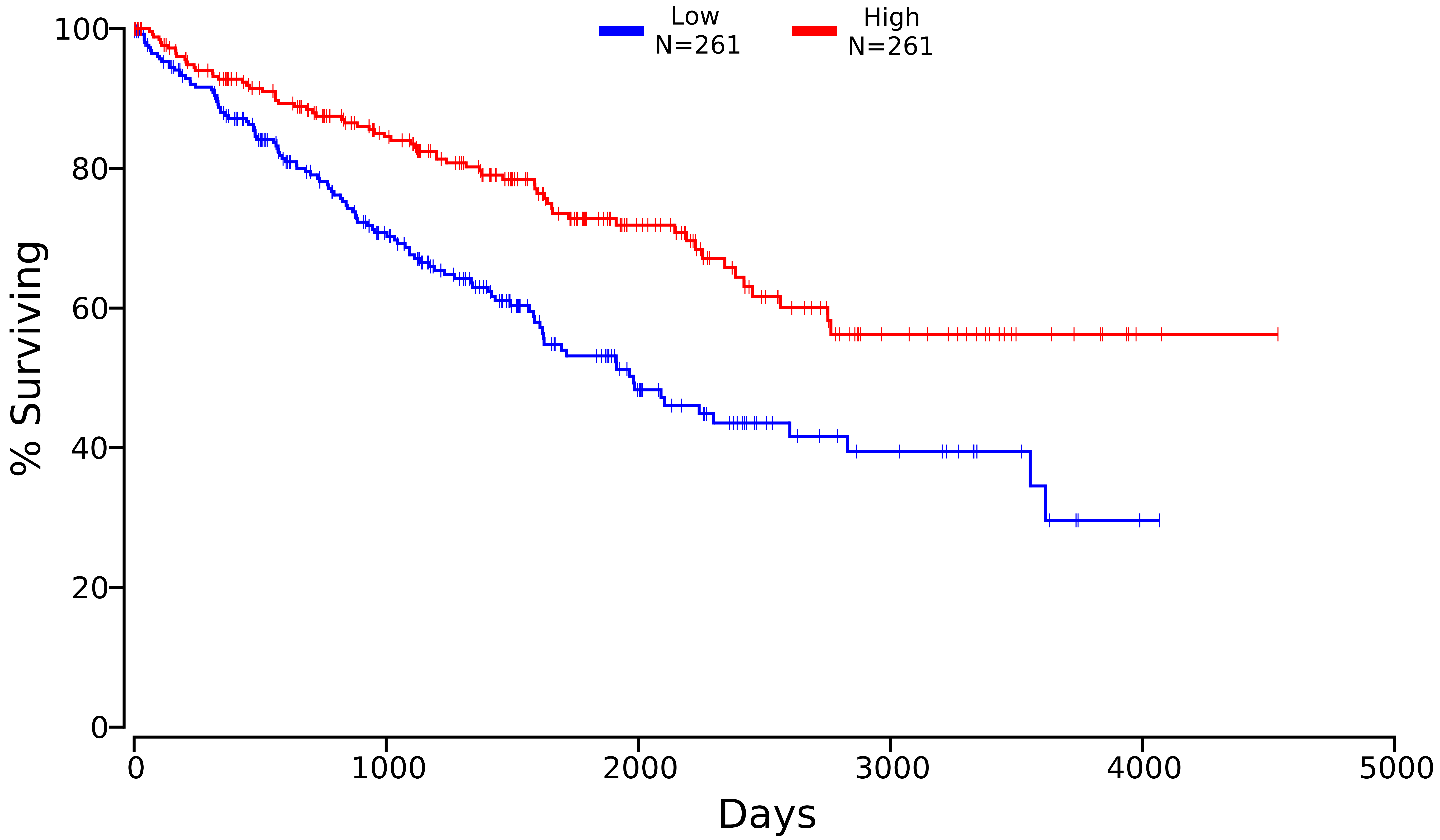

### PLVAP_KIRC_83483_50_50.pdf

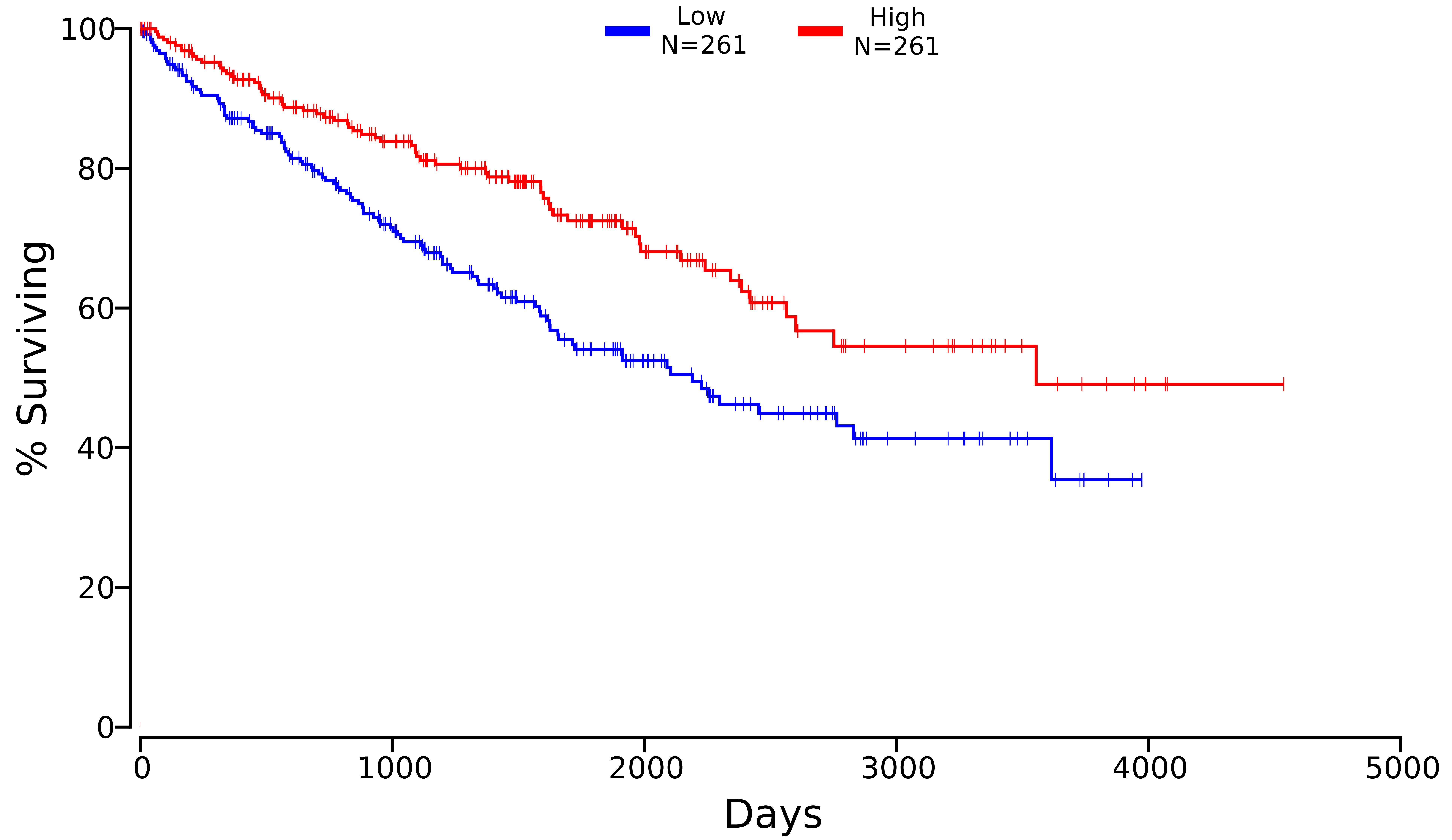

### PODXL_KIRC_5420_50_50.pdf

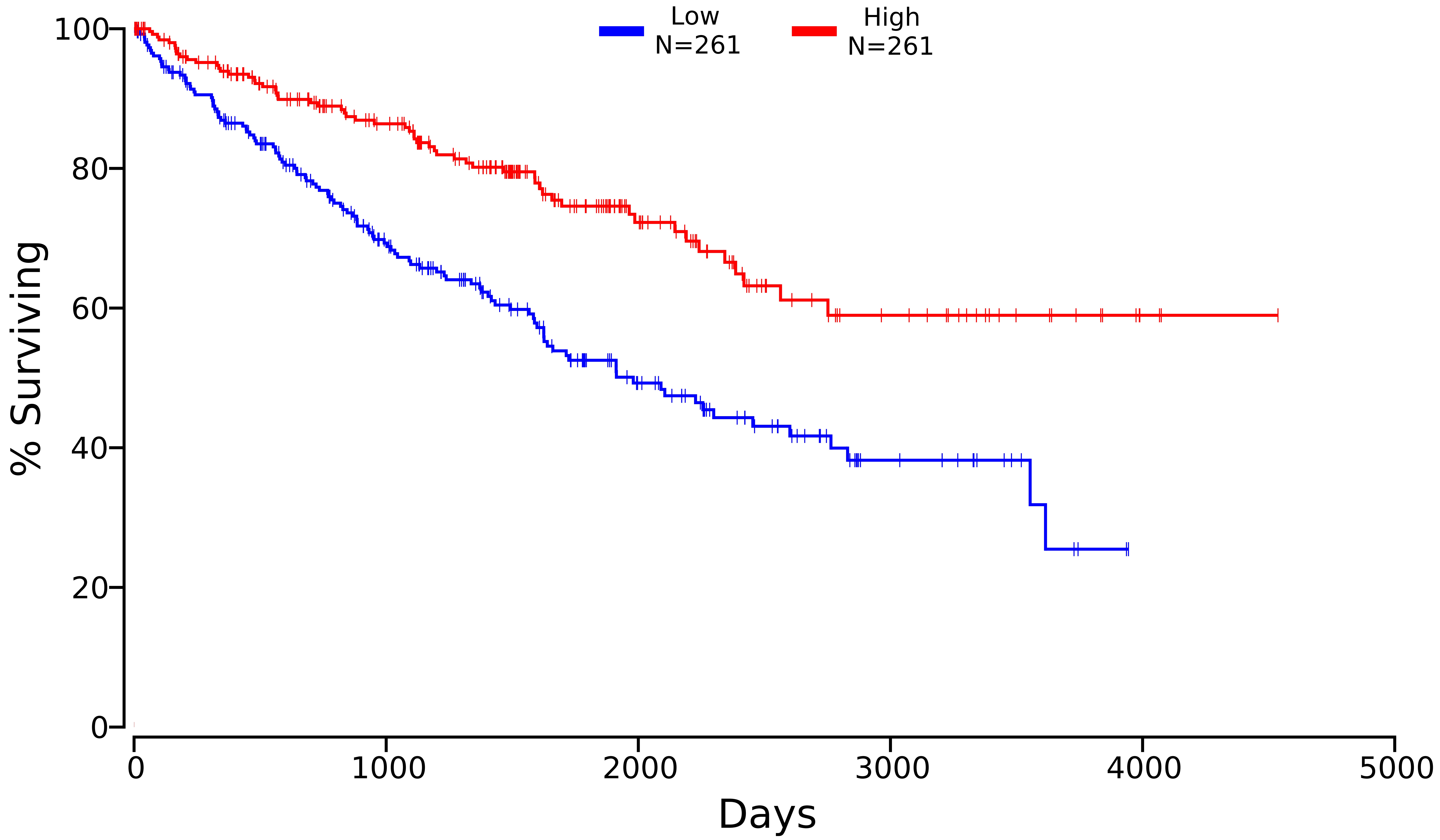

### TMBIM6_KIRC_7009_40_60.pdf

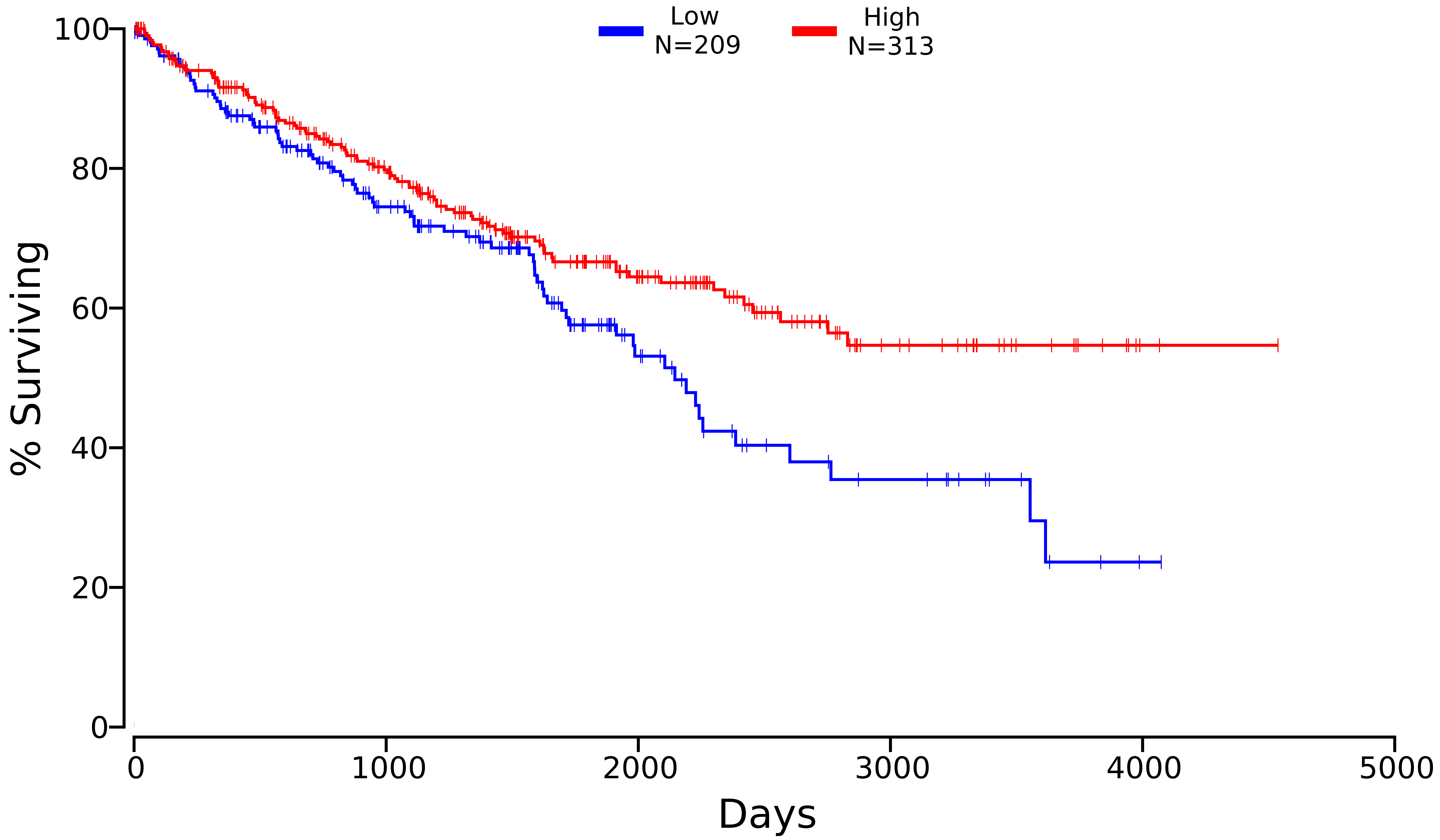

### VEGFA_KIRC_7422_70_30.pdf

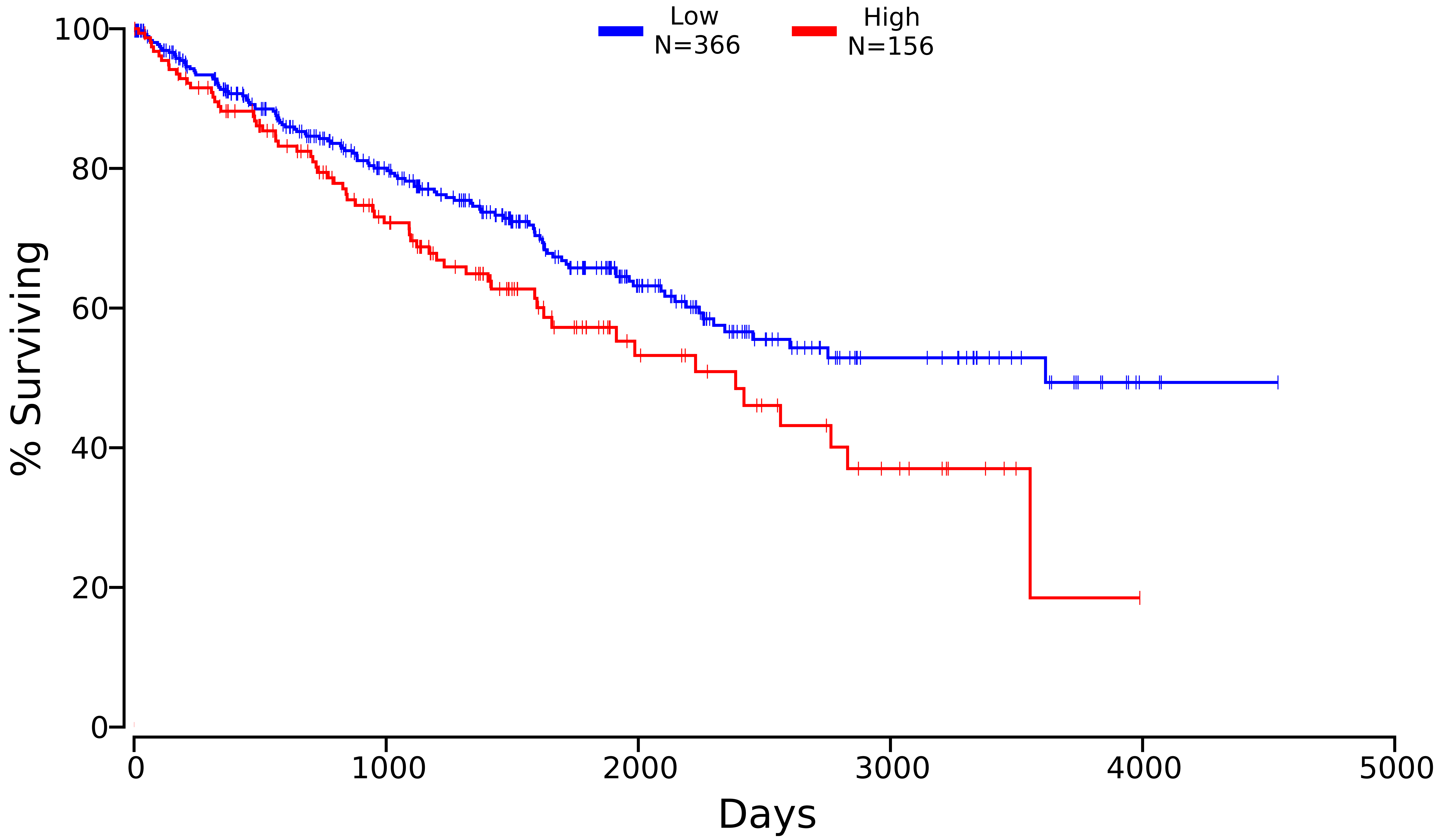

### VWF_KIRC_7450_50_50.pdf

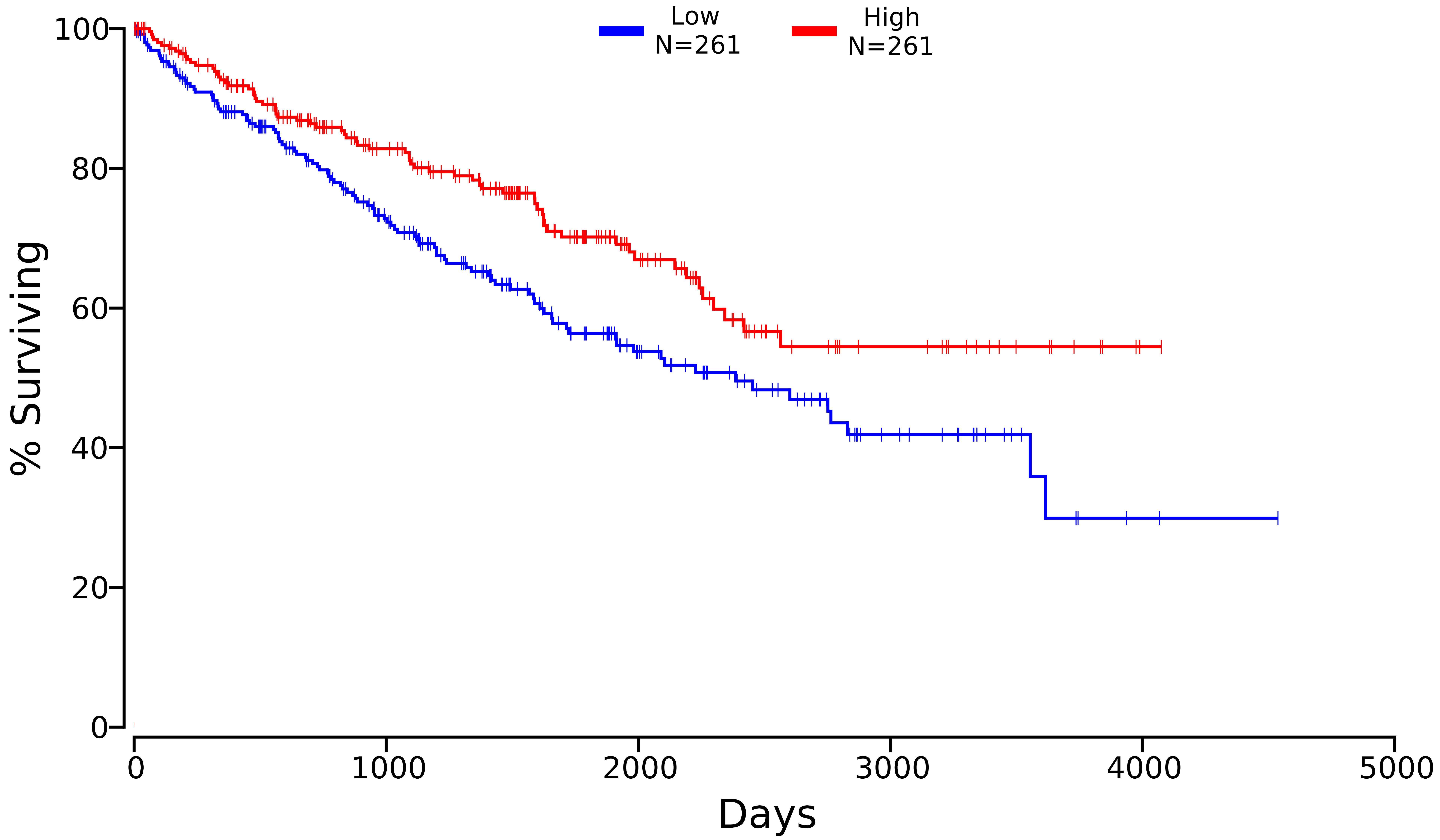
